# Evolutionarily labile pachytene piRNAs target an altered set of mRNAs in male hybrids of house mouse subspecies

**DOI:** 10.64898/2026.07.08.737336

**Authors:** Martin Säflund, Masomeh Askari, Atiyeh Eghbali, Mukhtar Mohamed Abdi, John L. Fitzpatrick, Tianxiong Yu, Deniz M. Özata

## Abstract

During male meiosis-I of placental mammals, ∼30-nucleotide pachytene PIWI-interacting RNAs (piRNAs) are expressed to regulate genes required for sperm function. Pachytene piRNA genes evolve rapidly. Whether rapid evolutionary turnover of pachytene piRNAs is under positive selective pressure remains enigmatic. Here, we investigate the evolutionary rate of pachytene piRNA genes over a short evolutionary timescale using geographically isolated mouse subspecies. We demarcate the genes producing postnatal piRNAs in PWK/PhJ and CAST/EiJ. Comparative genomics reveals 16 subspecies-specific pachytene piRNA loci underscoring how labile pachytene piRNA genes are even during short evolutionary timescale. We report a highly abundant CAST/EiJ-specific *pi17-CAST* locus defying the notion that young pachytene piRNA genes do not produce abundant piRNAs. In fact, male hybrids from the reciprocal crossing C57BL/6J and CAST/EiJ produce *pi17-CAST* piRNAs almost exclusively from the CAST/EiJ allele suggesting that species-specific nucleotide variants are sufficient to turn a locus into piRNA source. Intriguingly, hybrid males with reduced fertility features retain distinct piRNA-mRNA pairs compared to parents. Our work reveals that rapidly evolving pachytene piRNAs can gain or lose targets in the hybrid males of closely related mammalian species.

## Main

In animals, 21–35-nucleotide (nt) PIWI-interacting RNAs (piRNAs) associate with PIWI proteins and guide them to silence transposons, preserve genome integrity, and regulate mRNAs required for fertility^1–10^. Embryonic male germ cells produce fetal piRNAs that silence active transposons^1,2,8^. Shortly after birth, pre-pachytene piRNAs, which are ∼26-nt in length with unknown function, are produced mostly from the 3’ untranslated regions (3’ UTR) of mRNAs^11–13^. At the onset of the pachytene stage of male meiosis-I, ∼100 transposon-depleted loci (i.e., pachytene piRNA genes) generate long noncoding transcripts that are processed into highly abundant ∼30-nt pachytene piRNAs^11,12,14^. Timely production of pachytene piRNAs is collaboratively controlled by the transcription factors A-MYB and TCFL5^11,12,15–17^. In fact, A-MYB/TCFL5 regulatory axis drives the transcription of pachytene piRNA genes as well as genes encoding proteins that participate in piRNA biogenesis^11,12,16,17^.

Pachytene piRNA genes are required for reproduction; deletion of six major mouse pachytene piRNA genes disrupts male fertility^4,5,10,18^. Puzzlingly, pachytene piRNA genes diverge as rapidly as non-transcribed regions of the genome^12^. Rapid evolution of genes related to reproduction can have important consequences for generating reproductive isolation, thereby potentially driving speciation^19–22^. However, the lack of experimental mammalian system has obscured our understanding of the evolutionary function of rapidly evolving pachytene piRNAs. To date, in mice, pachytene piRNA genes are mapped in a single subspecies of house mice C57BL/6J (*Mus musculus domesticus*)^11^ that diverged from its siblings ∼0.5 million years ago^23–25^. Other wild-derived outbred house mice subspecies with distinct demographic and ecological histories provide a powerful yet under-explored system for studying the evolutionary function of pachytene piRNAs.

Here, we integrate piRNA-seq, RNA-seq, PRO-seq, CUT&RUN for histone H3 lysine 4 trimethylation (H3K4me3), and PAS-seq to demarcate the precise genomic coordinates of pachytene piRNA genes in PWK/PhJ (*Mus musculus musculus*) and CAST/EiJ (*Mus musculus castaneus*). As in C57BL/6J, 105 and 99 transposon-depleted genes, whose transcription is controlled by A-MYB/TCFL5 regulatory axis, produce pachytene piRNAs in PWK/PhJ and CAST/EiJ, respectively. Comparative genomics across three subspecies shows 16 subspecies-specific loci, demonstrating how rapidly pachytene piRNA genes evolve during short evolutionary timescale. Although most young loci produce fewer piRNAs compared with older loci, we unexpectedly map the CAST/EiJ-specific pachytene piRNA gene, *pi17-CAST*, which produces abundant piRNAs with notable number of putative target transcripts. Intriguingly, in hybrid males, *pi17-CAST* locus produced piRNAs almost exclusively from the CAST/EiJ allele suggesting that nucleotide variations acquired after the divergence of C57BL/6J and CAST/EiJ from their common ancestor are sufficient to turn the locus into a piRNA source. Furthermore, F2 hybrid males from reciprocal crossing of C57BL/6J and CAST/EiJ showed reduced fertility features compared to F1 hybrid males, regardless of the directionality of crossing. Notably, F2 hybrid males retain distinct piRNA-mRNA pairs compared to F1 hybrid males. Those mRNAs that are cleaved by piRNAs of F2 hybrid males with reduced fertility features were enriched for gene categories including sperm development and function. Our work thus provides evidence for the hitherto unrecognized role for pachytene piRNAs in building reproductive isolation.

## Results

### Two major classes of piRNA-producing loci in the genomes of wild-derived outbred mouse subspecies

To define loci producing piRNAs in the genomes of PWK/PhJ and CAST/EiJ, we sequenced piRNAs from seven developmental stages of testes (9 days postpartum [dpp], 11 dpp, 14 dpp, 17 dpp, 21 dpp, and 45 dpp) (Fig. 1a). Note that we pretreated small RNA samples with sodium periodate that oxidizes the 3’ ends of miRNAs and siRNAs and excludes them from sequencing libraries^12,26,27^. piRNAs resist oxidation because they are 2’-*O*-methylated at their 3’ ends in animals^27–32^. Using 5 kb sliding windows, we identified genomic regions containing ≥10 piRNAs/kb. Regions within 20 kb of each other were then merged into a piRNA locus with sufficiently high piRNA abundance (≥100 RPM; reads per million). This analysis identified 298 and 198 piRNA-producing loci in PWK/PhJ and CAST/EiJ, respectively (Fig. 1b, c; Supplementary Table 1a, b). Our defined piRNA-producing loci accounted for ∼95% of all piRNAs in the testes of PWK/PhJ and CAST/EiJ at 45 dpp (Extended Data Fig. 1a).

**Figure. 1.**
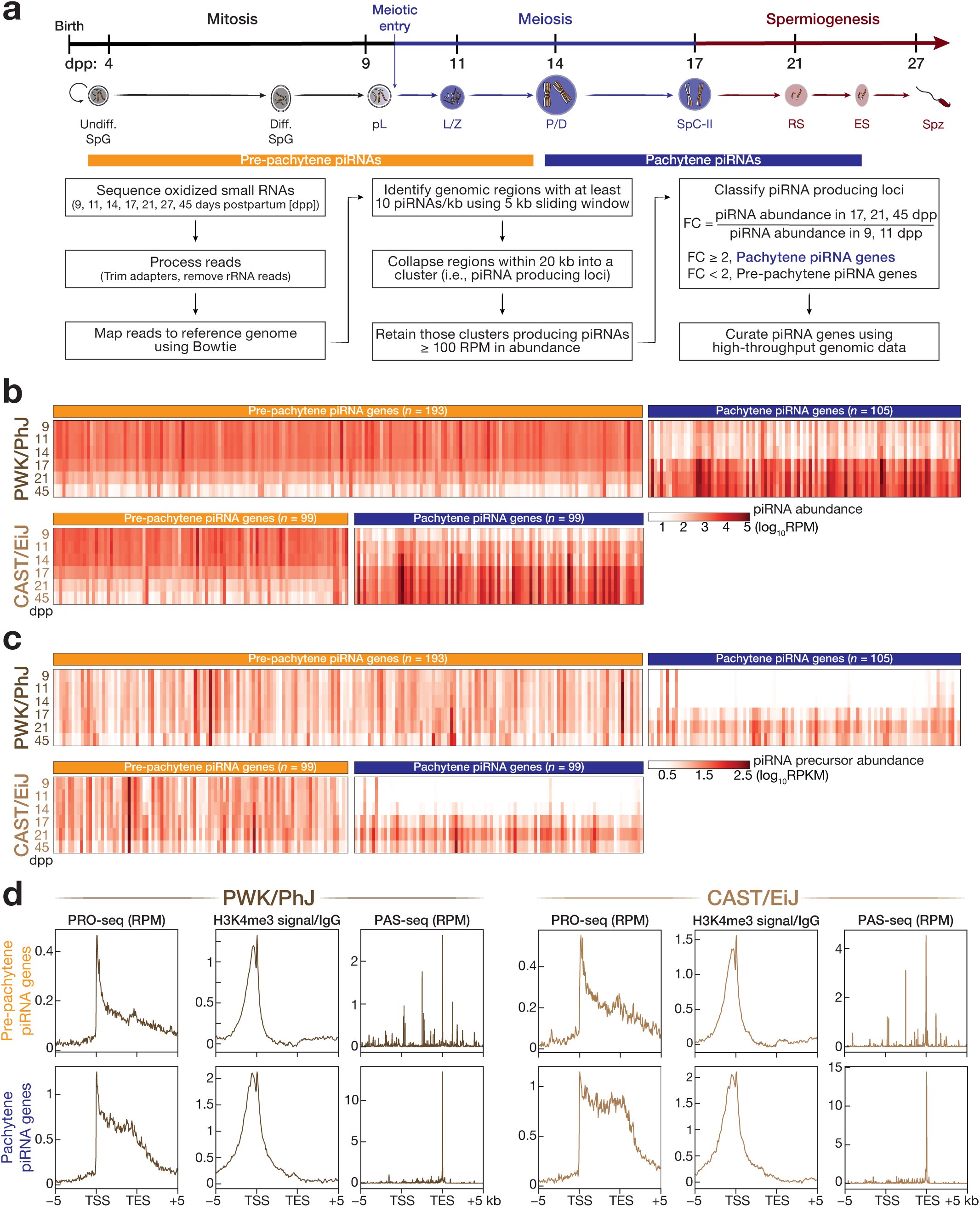
Testes of wild-derived outbred house mice subspecies express two classes of postnatal piRNA genes. **(a)** Schematic representation (top) for the developmental trajectory of male germ cells aligned with the age of mouse, days postpartum (dpp)^75^. Strategy (bottom) for defining postnatal piRNA genes in mice. Undiff. SpG, undifferentiated spermatogonia; Diff. SpG, differentiating spermatogonia; pL, preleptotene; L/Z, leptotene and zygotene spermatocytes; P/D pachytene and diplotene spermatocytes; SpC-II, secondary spermatocyte; RS, round spermatid; ES, elongating spermatid; Spz, spermatozoa. **(b)** Heatmaps show the abundance of reads per million (RPM)-normalized mature piRNAs produced from piRNA genes in the testes of PWK/PhJ (top) and CAST/EiJ (bottom) at different developmental stages (9, 11, 14, 17, 21, and 45 dpp). **(c)** Heatmap representation of the abundance of reads per kilobase of transcript per million (RPKM)-normalized piRNA precursor transcripts from piRNA genes across the testes of staged mice (9, 11, 14, 17, 21 and 45 dpp). PWK/PhJ (top) and CAST/EiJ (bottom). **(d)** Metaplots show RPM-normalized PRO-seq signal, fold enrichment of H3K4me3 CUT&RUN signal over IgG, and RPM-normalized PAS-seq in the –5 kb to +5 kb window flanking transcription start sites (TSSs) and transcription end sites (TESs) of pre-pachytene (top) and pachytene (bottom) piRNA genes in the genomes of PWK/PhJ (left) and CAST/EiJ (right).

Staged mouse testes reflect the temporal expression of the classes of postnatal piRNAs in C57BL/6J^11^: at 11 dpp, germ cells do not progress further than zygotene spermatocytes, and predominately express 26-27-nt pre-pachytene piRNAs, whereas by 17 dpp, when spermatogenesis reaches to late pachytene stage, expression of pachytene piRNAs peak^11^. To systematically classify piRNA-producing loci, we thus leveraged the temporal expression of piRNAs throughout the developmental stages of testes (Fig. 1a). Comparing the abundance of piRNAs from each piRNA-producing locus in 17 dpp, 21 dpp, and 45 dpp testes to that in 9 dpp and 11 dpp testes revealed two classes of piRNA-producing loci (Fig. 1b): (1) 193 PWK/PhJ and 99 CAST/EiJ loci, whose piRNA abundance changed <2-fold between 17–45 dpp testes and 9–11 dpp testes, were classified as pre-pachytene piRNA genes; (2) On the other hand, the abundance of piRNAs from 105 PWK/PhJ and 99 CAST/EiJ loci were ≥2-fold higher in 17–45 dpp testes than in 9–11 dpp testes. We thus classified those loci as pachytene piRNA genes (Fig. 1b; Supplementary Table 1a, b).

At 9–11 dpp, pre-pachytene piRNA genes dominated piRNA production. Here, >90% of all piRNAs mapped to pre-pachytene piRNA genes (PWK/PhJ, median pre-pachytene piRNA abundance = 902 RPM vs. median pachytene piRNA abundance = 64 RPM; CAST/EiJ, median pre-pachytene piRNA abundance = 1,689 RPM vs. median pachytene piRNA abundance = 118 RPM; Fig. 1b; Supplementary Table 2a, b). However, at 17–45 dpp, pachytene piRNAs accounted for >87% of all piRNAs. In fact, the median abundance of piRNAs mapping to pachytene piRNA genes increased 30-and 15-fold in 17–45 dpp testes when compared with 9–11 dpp testes in PWK/PhJ and CAST/EiJ, respectively (Fig. 1b; Supplementary Table 2a, b). Furthermore, the abundance of piRNA precursor transcripts corroborated our gene classification: The transcript abundance from pre-pachytene piRNA genes remained nearly constant between 9-11 dpp testes (PWK/PhJ, median = 9.1 reads per kilobase of transcript per million normalized (RPKM); CAST/EiJ, median = 9.5 RPKM) and 17–45 dpp testes (PWK/PhJ, median = 8.3 RPKM; CAST/EiJ, median = 11.2 RPKM). By contrast, the abundance of piRNA precursor transcripts from pachytene piRNA genes was >300-fold higher in 17–45 dpp than in 9–11 dpp testes in both PWK/PhJ and CAST/EiJ (Fig. 1c; Supplementary Table 2c, d).

The majority of the pre-pachytene piRNA genes corresponded to protein-coding genes (191 of 193 PWK/PhJ genes; 95 of 99 CAST/EiJ genes; exemplified in Extended Data Fig. 1b). However, most of the pachytene piRNA genes resided within regions that do not encode proteins, consistent with the typical characteristics of pachytene piRNA genes in C57BL/6J (70 of 105 PWK/PhJ genes; 74 of 99 CAST/EiJ genes; exemplified in Extended Data Fig. 1c). As in C57BL/6J, ∼50% of the pachytene piRNA genes were intronless, whereas only ∼3% of pre-pachytene piRNA genes lacked introns consistent with the fact that pre-pachytene piRNAs are derived from protein-coding genes (Extended Data Fig. 2a). Those pachytene piRNA precursor transcripts retaining introns were spliced prior to processing into piRNAs: median piRNA abundance from the exons of pachytene piRNA genes (PWK/PhJ, median = 1,862 RPM; CAST/EiJ, median = 2,042 RPM) was >500-fold greater than from the introns (PWK/PhJ, median = 3.7 RPM; CAST/EiJ, median = 3.8 RPM; Extended Data Fig. 2a). We thus conclude that in wild-derived outbred mice, pachytene piRNA precursor transcripts are transcribed from discrete genomic loci and are spliced before being processed into piRNAs at the pachytene stage of male meiosis-I.

### Transposon-depleted pachytene piRNA genes are transcribed by Pol II in wild-derived outbred house mice subspecies

We performed PRO-seq, CUT&RUN for H3K4me3, and PAS-seq from FACS-purified pachytene and diplotene (P/D) spermatocytes of PWK/PhJ and CAST/EiJ to curate both the transcription start sites (TSSs) and the transcription end sites (TESs) of our annotations (Fig. 1d). PRO-seq measures the distribution of transcriptionally engaged RNA polymerase II (Pol II) over genes^33^. Typically, Pol II peak accumulates around the TSSs of genes representing promoter-proximal pausing, whilst Pol II pausing serves as an important regulatory step in coordinating transcriptional elongation. For example, Pol II pausing coincides with the 5’end capping of transcripts^33–37^. Our PRO-seq data revealed prominent Pol II peaks near the TSSs of piRNA genes that were annotated in P/D spermatocytes of PWK/PhJ and CAST/EiJ mice (Fig. 1d). CUT&RUN for H3K4me3 – a histone modification at promoter-proximal regions of genes transcribed by Pol II^38^ – from the P/D spermatocytes of PWK/PhJ and CAST/EiJ mice further confirmed the TSS of each piRNA gene (Fig. 1d). Whereas our PAS-seq data captured the 3’ ends of piRNA precursor transcripts that precede their poly(A) tails (Fig. 1d).

Both in humans and C57BL/6J mice, pachytene piRNA genes contain fewer transposons than the whole genome^11,12,14,39,40^. Consistently, transposon expression remained unchanged in C57BL/6J males lacking *pi6*^10^ or *pi18*^5^, attributing to the function of pachytene piRNAs beyond transposon silencing. Like C57BL/6J, the transposon content of the exons of pre-pachytene piRNA precursors from wild-derived outbred mice (PWK/PhJ, median = 15.1; CAST/EiJ, median = 14.6) were similar to that of mRNAs that are not source of piRNAs (median = 18.4) (Extended Data Fig. 2b). Even though pachytene piRNA precursors are long non-coding RNAs, their transposon content within exons (PWK/PhJ, median = 28.6; CAST/EiJ, median = 32.2) were less than the exons of other lncRNAs which do not make piRNAs (median = 42.8), suggesting a purifying selection on transposons within the exons of pachytene piRNA genes (Extended Data Fig. 2b). We conclude that, as in C57BL/6J, pachytene piRNA precursor transcripts are transcribed by Pol II and are devoid of transposon sequences.

### A-MYB/TCFL5 regulatory axis regulates pachytene piRNA production in wild-derived outbred mouse subspecies

In C57BL/6J mice, the transcription factor (TF) A-MYB initiates the transcription of TF *Tcfl5*, whose protein product subsequently regulates both its own and *A-Myb* expression, establishing a mutually reinforcing positive feedback circuit at the pachytene stage of male meiosis-I^17,41^. To test whether this positive feedback loop between A-MYB and TCFL5 is conserved in mouse subspecies, we performed CUT&RUN for A-MYB and TCFL5 from FACS-purified P/D spermatocytes of PWK/PhJ and CAST/EiJ (Extended Data Fig. 3). Here, we detected A-MYB occupancy around the TSSs of both *A-Myb* and *Tcfl5* genes (Extended Data Fig. 3a). Similarly, TCFL5 CUT&RUN located a prominent TCFL5 peak near the TSSs of *A-Myb* and *Tcfl5* in P/D of PWK/PhJ and CAST/EiJ (Extended Data Fig. 3a).

The A-MYB/TCFL5 regulatory axis triggers pachytene piRNA production by activating the transcription of pachytene piRNA genes, as well as genes encoding piRNA biogenesis proteins in C57BL/6J mice^11,12,17^. To test if this circuit regulates piRNA production in outbred mouse species, we determined the genes whose promoter-proximal regions are bound by A-MYB/TCFL5 regulatory axis. To this end, we computed the distance from the TSSs of pachytene piRNA and piRNA biogenesis genes to the nearest significant A-MYB or TCFL5 peaks. Those with both A-MYB and TCFL5 peaks or with either A-MYB or TCFL5 peak within ±2 kb of their TSSs were considered as genes regulated by A-MYB/TCFL5 regulatory axis. CUT&RUN data demonstrated that >90% of pachytene piRNA genes were under the control of the A-MYB/TCFL5 regulatory axis (Extended Data Fig. 3b; Supplementary Table 3). The promoters of 89 of the 105 PWK/PhJ and 91 of the 99 CAST/EiJ pachytene piRNA genes were occupied by both A-MYB and TCFL5, while the promoters of seven and six pachytene piRNA genes were bound by either of the factors in PWK/PhJ and CAST/EiJ, respectively (Extended Data Fig. 3b; Supplementary Table 3). Furthermore, the promoters of all 20 piRNA biogenesis genes were bound by TCFL5 (Extended Data Fig. 3b; Supplementary Table 3; PWK/PhJ, median distance = 112 bp; CAST/EiJ, median distance = 170 bp). Similarly, 19 of the 20 piRNA biogenesis genes featured A-MYB peak near their TSSs (Extended Data Fig. 3b; Supplementary Table 3; PWK/PhJ, median distance = 132 bp; CAST/EiJ, median distance = 105 bp). Finally, exemplified two pachytene piRNA and six piRNA biogenesis genes corroborated our findings: the promoters of pachytene piRNA and piRNA biogenesis genes exhibited significant A-MYB and TCFL5 peaks near their TSSs (Extended Data Fig. 3c, d). We conclude that, like in C57BL/6J, pachytene piRNA production is regulated by A-MYB/TCFL5 regulatory axis in PWK/PhJ and CAST/EiJ.

### Several pachytene piRNA genes lacked conservation across house mouse subspecies

Next, comparing the genomic coordinate (i.e., synteny) and piRNA abundance of each pachytene piRNA gene we demarcated across the three house mice subspecies revealed that 88 genes produced pachytene piRNAs at a density of ≥100 RPM from the syntenic location in all three subspecies – i.e. conserved pachytene piRNA genes. This suggested that many pachytene piRNA genes antedate the divergence of the three subspecies from their last common ancestor (Fig. 2a). Of note, these 88 conserved pachytene piRNA loci included one additional pachytene piRNA gene that was not annotated in the earlier C57BL/6J annotation^11^. Conversely, 16 pachytene piRNA genes did not produce sufficient piRNAs (i.e. non-conserved pachytene piRNA genes) in one or two of the three subspecies despite the conservation of their genomic coordinates, implying that these loci were no older than ∼0.5 million years^23–25^. Of the 16 non-conserved genes, nine were found in a maximum two subspecies. The remaining seven loci produced piRNAs in only one of the subspecies (Fig. 2a). Moreover, we identified 11 conserved loci that produced different classes of piRNAs across the three subspecies. Of these, all were classified as pachytene piRNA genes in PWK/PhJ, whereas seven and 10 were a source for other classes of piRNAs in the genomes of CAST/EiJ and C57BL/6J, respectively (Fig. 2a; Supplementary Table 2).

**Figure. 2.**
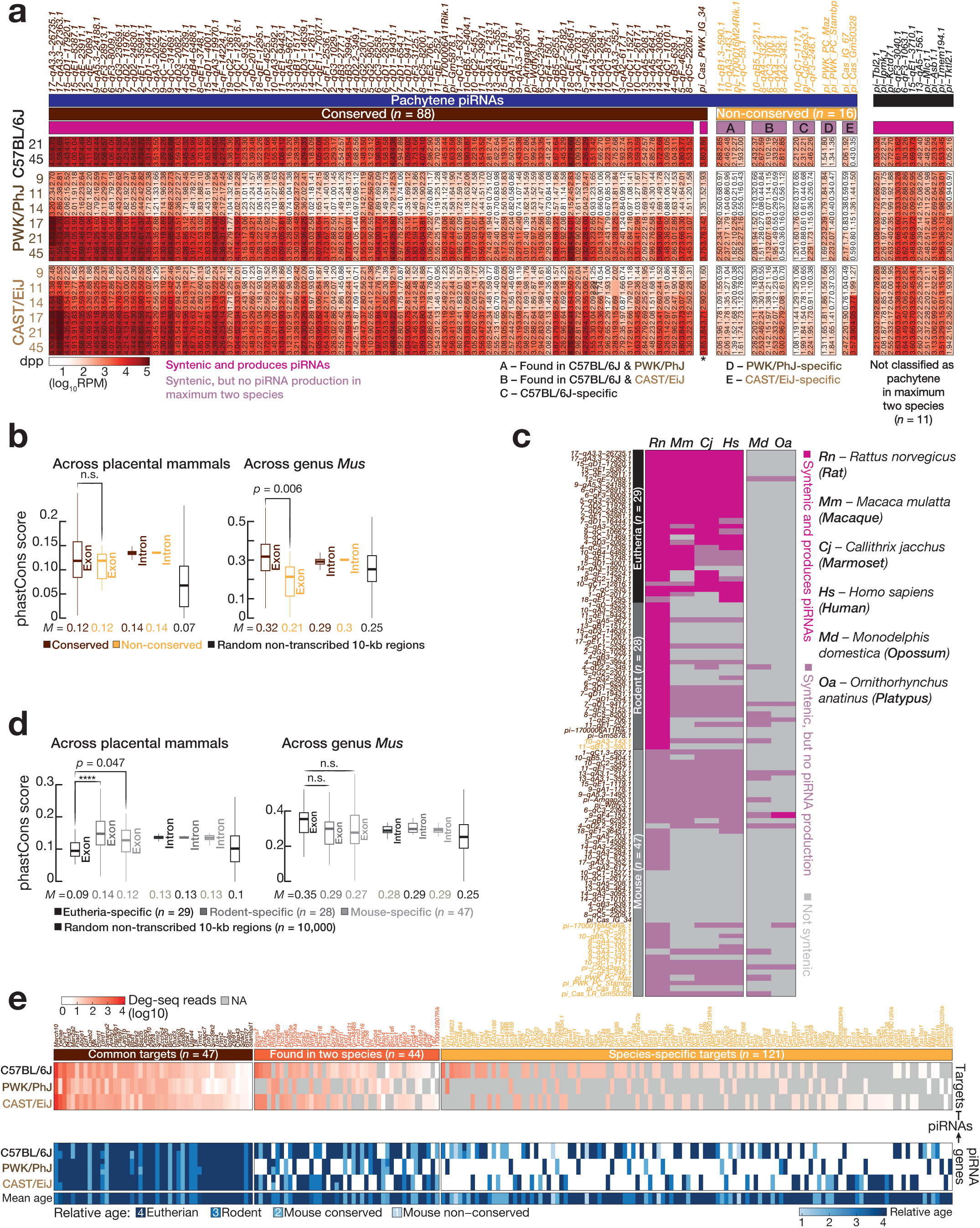
Mouse pachytene piRNA genes rapidly evolve. (a) Each column represents a pachytene piRNA gene grouped as conserved or non-conserved genes by its function as relative piRNA abundance and its genomic conservation across mice subspecies. piRNA abundance is reported as log_10_ transformed reads per million (RPM) with a pseudo-count of 1. (b) Boxplots show DNA sequence conservation for the exons and introns of conserved and non-conserved mouse pachytene piRNA genes across placental mammals (left) or genus *Mus* (right). Randomly selected non-transcribed 10-kb regions served as background control. Horizontal lines represent the median. Whiskers depict the 95% confidence intervals. Interquartile range (IQR) is represented by boxplots. Significance is measured using two-sided unpaired Student *t* test. (c) Synteny analysis shows the conservation of genomic coordinates of mouse pachytene piRNA genes across placental and non-placental mammals. piRNA-producing loci from rat, macaque, marmoset, human, opossum and platypus were defined in our previous work^12^. (d) Boxplots show DNA sequence conservation for the exons and introns of eutherian, rodent-specific, and mouse-specific pachytene piRNA genes across placental mammals (left) and across genus *Mus* (right). Randomly selected non-transcribed 10-kb regions served as background control. Horizontal lines represent the median. Whiskers depict the 95% confidence intervals. Interquartile range (IQR) is represented by boxplots. Significance is measured using two-sided unpaired Student *t* test. (e) Heatmap (top) shows the log_10_ transformed abundance of piRNA-guided cleaved transcripts across mice subspecies. Heatmap (bottom) shows the relative evolutionary age of pachytene piRNA genes whose piRNAs direct the cleavage of transcripts.

### Pachytene piRNA genes are highly labile in the genomes of placental mammals

Pachytene piRNA genes are poorly conserved across species^12,42,43^. Indeed, DNA sequences of human pachytene piRNA genes evolve as fast as non-transcribed regions in the genomes of placental mammals^12^ (∼160 million years^25,44^). As such, we then sought to examine the sequence divergence rate for mouse pachytene piRNA genes across other placental mammals. As expected, the coding sequences of both protein-coding genes that do not produce piRNAs, and protein coding genes, which make pre-pachytene piRNAs, were the most conserved transcribed features in the genomes of 40 placental mammals (Extended Data Fig. 4a; protein coding genes not making piRNAs, median phastCons score = 0.75; pre-pachytene piRNA genes, median phastCons score = 0.88), suggesting that coding sequences of protein coding genes are under purifying selection. Although the DNA sequences of promoter-proximal regions of pachytene piRNA genes were conserved (Extended Data Fig. 4a; right), the transcribed regions of pachytene piRNA genes lacked conservation. In fact, the exons of pachytene piRNA genes, from which piRNAs derive, diverged at a similar rate as that of lincRNAs and randomly selected non-transcribed regions (Extended Data Fig. 4a; left). We next investigated how rapidly mouse pachytene piRNA genes evolve over shorter evolutionary timescales in the genomes of 19 species from the genus *Mus* (∼7 million years^25,44^). Here, the exons of pachytene piRNA genes were among the least conserved genomic features despite the conservation of their promoters in the genomes of 19 mouse species (Extended Data Fig. 4b).

We found that several pachytene piRNA genes are specific to one or at most two mice subspecies which diverged from their last common ancestor ∼0.5 million years ago ^23–25^, suggesting that pachytene piRNA genes are among the most labile transcription units within the genomes of placental mammals (i.e., non-conserved genes; Fig. 2a).

This prompted us to investigate whether conserved and non-conserved pachytene piRNA genes are under a different evolutionary force. Here, the exons of non-conserved pachytene piRNA genes diverged at a similar rate as that of conserved pachytene piRNA genes across placental mammals (Fig. 2b; left). Notably however, in the genomes of the genus *Mus*, DNA sequences of exons from non-conserved genes diverged faster compared to that of conserved genes, while the introns diverged at a similar rate (Fig. 2b; right; two-sided unpaired Student *t* test; conserved genes, median phastCons score = 0.32; non-conserved genes, median phastCons score = 0.21). To further substantiate this observation, we examined the conservation of genomic coordinates for mouse pachytene piRNA genes across other mammals using our previously annotated piRNA-producing loci in macaque, marmoset, rat, opossum – a marsupial, and platypus – a monotreme^12^ (Fig. 2c; Supplementary Table 4). Among the 104 mouse pachytene piRNA genes, 29 produced piRNAs from the syntenic location in at least two other placental mammals (i.e. eutherian genes). Another 28 loci produced piRNAs only in rats other than mice (i.e. rodent-specific), while the remaining 47 loci were specific to mouse (i.e. mouse-specific) (Fig. 2c; Supplementary Table 4). Of note, two of the non-conserved loci (Fig. 2a; *10–qA3–143.1* and *11–qB1.3–590.1*) were found in rats suggesting that they were lost in CAST/EiJ. Henceforth, mouse- and rodent-specific loci were named young pachytene piRNA genes, whereas eutherian loci were termed old pachytene piRNA genes. Nonetheless, only one of the 104 loci was found in platypus suggesting that pachytene piRNAs are acquired in placental mammals after the divergence of placental and non-placental mammals (Fig. 2c; ∼160 million years^25,44^). We next examined the divergence rate for the exons of old and young pachytene piRNA genes during long (i.e. across placental mammals) and short (i.e. across genus *Mus*) evolutionary time scales (Fig. 2d). When we compared DNA sequence conservation among placental mammals, exons of old pachytene piRNA genes – that produce piRNA across placental mammals – evolved faster than that of young genes that are not piRNA source in species beyond rats (Fig. 2d). In contrast, we observed faster divergence of exons of young pachytene piRNA genes when compared with that of old genes in the genomes of genus *Mus* despite the lack of statistical significance (Fig. 2d). Taken together, our data suggests that once a locus becomes a piRNA source across a clade or within an order/genus, divergence of its DNA sequence is accelerated, implying increased sequence variation is related to piRNA function.

### Species-specific piRNA-guided cleaved transcripts appear across house mouse subspecies

The majority of C57BL/6J pachytene piRNAs have no target transcripts^4,10^, yet counterintuitively, several pachytene piRNA genes are required for functional sperm^4,5,10^. We found that mouse pachytene piRNA genes are highly labile across placental mammals, even among genus *Mus*. To begin to understand the biological meaning of the high evolutionary drift of pachytene piRNAs across mice subspecies, we sequenced 5’-monophosphorylated long RNAs (degradome-seq) from the 45 dpp testes of C57BL/6J, PWK/PhJ, and CAST/EiJ. piRNAs guide PIWI proteins to slice target transcripts, resulting in 5’-monophosphorylated 3’cleavage products^4,10,18,45^. We complemented our degradome-seq data with piRNA sequencing data to identify putative piRNA-guided 3’cleavage products (Supplementary Table 5). To be qualified as a putative target, we searched for guide nucleotides g2 to g15 with perfect complementarity between a piRNA and a cleaved RNA fragment (Methods). Our analysis identified 47 piRNA-guided cleaved transcripts for which at least 20 degradome-seq reads were found in all three mice subspecies (i.e., common targets). Notably, we found 121 piRNA-target pairs that were captured only in one of the three subspecies (i.e. species-specific). Of the 121 species-specific targets, 58 were specific to C57BL/6J, whilst 20 and 43 were found in PWK/PhJ and CAST/EiJ, respectively (Fig. 2e; Supplementary Table 6). Acquisition of species-specific piRNA-target pairs after the divergence of house mice subspecies from their common ancestor (∼0.5 million years^23–25^) provides evidence for the idea that rapidly evolving piRNAs target distinct sets of transcripts during mammalian evolution^12^.

### CAST/EiJ-specific locus produces unusually abundant pachytene piRNAs with putative target transcripts

Typically, the depth of synteny of a pachytene piRNA locus predicts its piRNA abundance, meaning that older loci produce more piRNAs than younger genes^12,43^. Consistently, in all three mice subspecies, the median abundance of pachytene piRNAs from conserved genes (C57BL/6J, median = 3,467 RPM; PWK/PhJ, median = 2,436 RPM; CAST/EiJ, median = 2,797 RPM) was ≥20-fold greater than that of pachytene piRNAs from non-conserved genes (C57BL/6J, median = 174 RPM; PWK/PhJ, median = 75 RPM; CAST/EiJ, median = 90 RPM) (Fig. 3a; two-sided unpaired Student *t* test). Intriguingly, we discovered a CAST/EiJ-specific pachytene piRNA gene, *pi-Gm50328* (Henceforth, *pi17-CAST*), that produces highly abundant pachytene piRNAs. In fact, in CAST/EiJ, the abundance of pachytene piRNAs from *pi17-CAST* (piRNA abundance = 6,918 RPM) was ∼2.5-fold higher than the median abundance of pachytene piRNAs from conserved genes (Fig. 3a). In CAST/EiJ, *pi17-CAST* locus is transcribed from a 1,125-bp long, bidirectional promoter marked by H3K4me3. Here, piRNAs were produced on the positive strand, while the *Lemd2* mRNA was generated from the negative strand on the opposite arm (Fig. 3b). On the contrary, *pi17-CAST* locus was transcriptionally off in C57BL/6J and PWK/PhJ. Indeed, PRO-seq from P/D spermatocytes of C57BL/6J and PWK/PhJ revealed no active transcription from *pi17-CAST* locus despite the comparable transcription rate of *Lemd2* across subspecies (Fig. 3b).

**Figure. 3.**
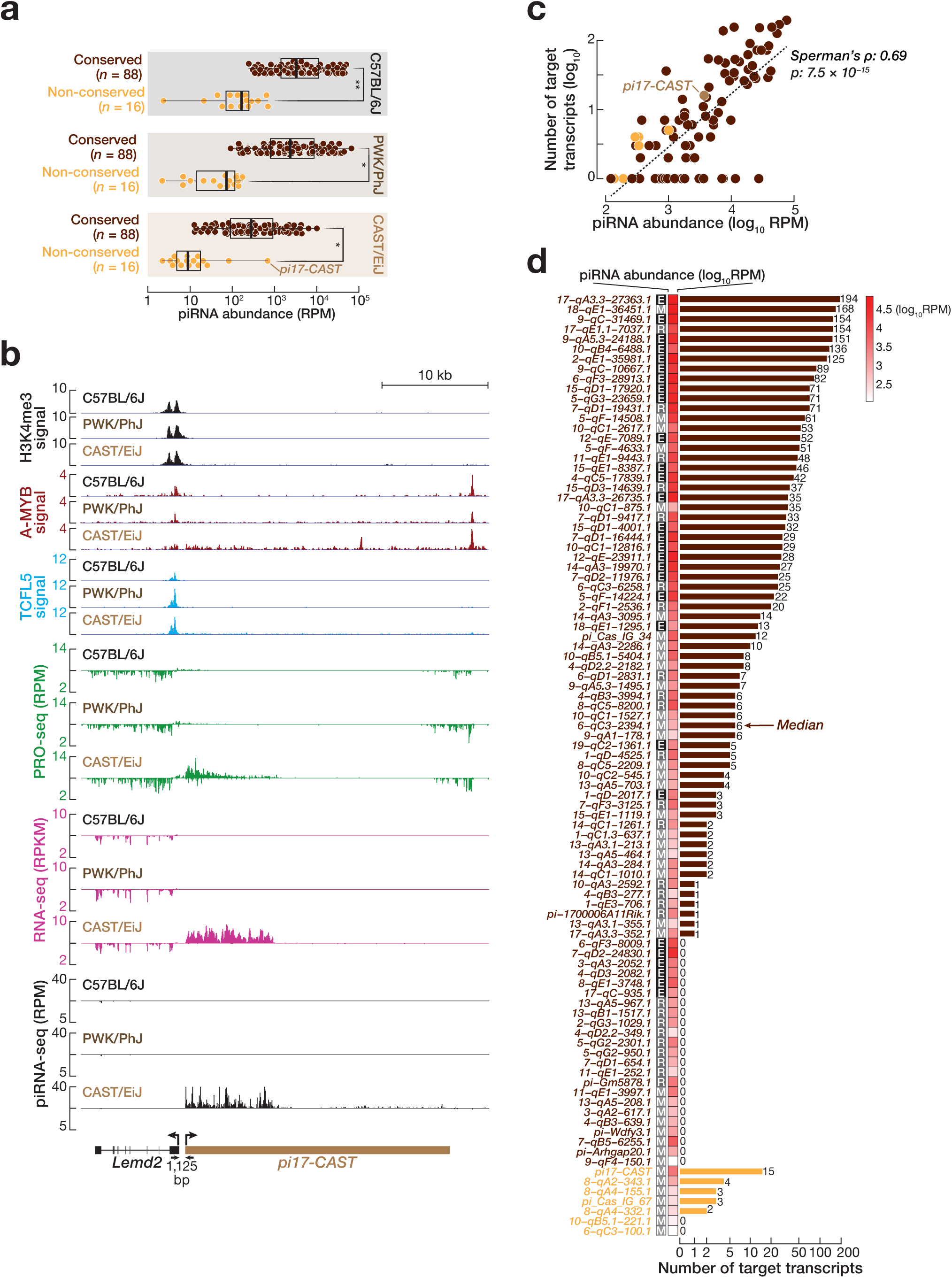
CAST/EiJ-specific *pi17-CAST* pachytene piRNA gene produces unusually abundant piRNAs. **(a)** Boxplots show the relative read per million (RPM)-normalized abundance of piRNAs from conserved and non-conserved pachytene piRNA genes in each subspecies. Each dot represents the average abundance of piRNAs from three biological replicates for each pachytene piRNA gene in adult testis (45 days postpartum [dpp]). Vertical lines represent the median. Whiskers depict the 95% confidence intervals. Interquartile range (IQR) is represented by boxplots. Significance is measured using two-sided unpaired Student *t* test. **(b)** Integrative Genomics Viewer (IGV) view of *pi17-CAST* locus across mice subspecies. H3K4me3, AMYB, and TCFL5 occupancies at promoter-proximal region. Signal tracks for H3K4me3 CUT&RUN, A-MYB and TCFL5 CUT&RUN, and PRO-seq are from FACS-purified pachytene/diplotene spermatocytes except for RNA-seq and piRNA-seq signal, which are from adult testis at 45 days postpartum. Signals for CUT&RUN, PRO-seq, and piRNA-seq are reported as reads per million (RPM). RNA-seq signal is reported as reads per kilobase of transcript per million (RPKM). **(c)** Scatter plot shows the spearman’s correlation (P) between the number of putative target transcripts and the abundance of piRNAs for each pachytene piRNA gene. Each dot represents the average of three biological replicates. **(d)** Bar plot represents the total number of putative target transcripts that are targeted by piRNAs from each pachytene piRNA gene. Of note, putative target transcripts exclude transposons.

The abundance of piRNAs is one of the key determinants of effective target silencing^18,46,47^. Consistently, pachytene piRNA abundance of a locus predicted its number of putative targets: for each locus, piRNA abundance and number of its targets were strongly correlated (Fig. 3c; Spearman’s ρ = 0.69; *p* = 7.5 × 10^−15^). Our degradome-seq identified 15 putative target transcripts cleaved by piRNAs from *pi17-CAST* that was more than twice the median number of target transcripts cleaved by piRNAs from each conserved piRNA locus (Fig. 3d; Supplementary Table 5c). Of the 15 putative targets, five were mRNAs (*Cdh24*, *Crebrf*, *Mtfr1*, *Knstrn*, *Gatad2a* and *Iba57*), while nine were other piRNA precursor transcripts (Extended Data Fig. 5; Supplementary Table 5c), supporting the general view^4,10,48^ that many pachytene piRNAs act ‘selfishl’ to amplify the abundance of other pachytene piRNAs. Taken together, we conclude that *pi17-CAST* produces highly abundant pachytene piRNAs with a notable number of putative target transcripts.

### Hybrid males produce *pi17-CAST* piRNAs from the CAST/EiJ allele

High prevalence of structural variations (SVs; ≥50 bp in size) within pachytene piRNA genes may partly explain the rapid divergence of pachytene piRNA genes across mammals^49^. However, the comparison of *pi17-CAST* locus across three subspecies found no SVs (Extended Data Fig. 6a).

We found that young pachytene piRNA genes appear to diverge faster when compared to old genes (Fig. 2), suggesting that accumulating mutations might suffice for turning a locus into a piRNA source. To test the idea that CAST/EiJ-specific single nucleotide polymorphisms (SNPs) underlie the piRNA production from *pi17-CAST* locus, we reciprocally crossed C57BL/6J and CAST/EiJ mice and sequenced piRNAs from the F1 hybrid males. The abundance of *pi17-CAST* piRNAs was nearly halved in hybrid males when compared with CAST/EiJ males (Fig. 4a, b; Extended Data Fig. 6b). Given that *pi17-CAST* locus produces no piRNAs in C57BL/6J, the ∼50% reduction in piRNA abundance implied that piRNAs are produced from the CAST/EiJ allele in F1 hybrids (Fig. 4b; Extended Data Fig. 6b). To test this, using piRNA reads, we first identified confident single nucleotide polymorphisms (SNPs) between C57BL/6J and CAST/EiJ and measured the abundance of piRNA reads mapping to those nucleotide variants in F1 hybrids, C57BL/6J, and CAST/EiJ mice. Two pachytene piRNA genes, *2–qE1–35981* (a eutherian; *pi2*) and *7–qD1–19431* (a rodent-specific; *pi7*), served as control. Intriguingly, in F1 hybrids, *pi17-CAST* piRNAs with subspecies-specific SNPs corresponded almost exclusively CAST/EiJ nucleotide variants, whereas *pi2* or *pi7* piRNAs with subspecies-specific SNPs were equally distributed between C57BL/6J and CAST/EiJ alleles (Fig. 4c; Extended Data Fig. 6c). We next calculated the frequency of each CAST/EiJ-specific SNP by computing the fraction of piRNA reads mapping to CAST/EiJ alleles. Here, the frequency of CAST/EiJ-specific SNPs in *pi17-CAST* gene, in the hybrid males, was almost 1 (Fig. 4d; median = 1; mean = 0.98–1; two-sided wilcoxon matched-pairs signed rank sum test, *p* values ≤ 1.3 × 10^-5^), whereas CAST/EiJ alleles in *pi2* or *pi7* genes were represented at ∼0.5 frequency in hybrid males (Fig. 4d; Extended Data Fig. 6d; *pi2,* median = 0.43-0.46, mean = 0.43-0.46; *pi7,* median = 0.47-0.52, mean = 0.46-0.50). We thus conclude that *pi17-CAST* locus lacking SVs produces piRNAs almost exclusively from the CAST/EiJ allele in hybrid males.

**Figure 4.**
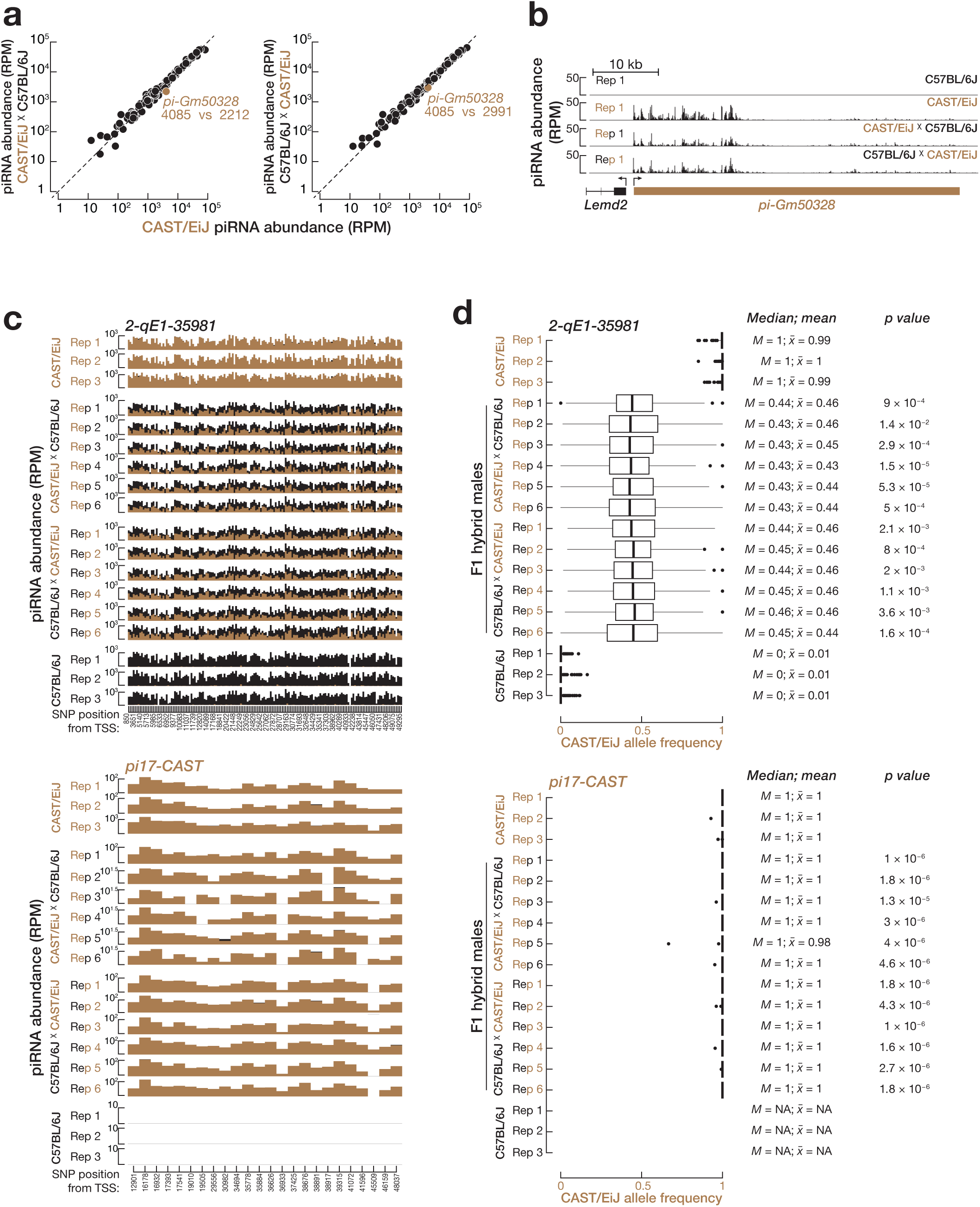
*pi17-CAST* produces piRNAs from CAST/EiJ allele in F1 hybrid males. **(a)** Scatter plots show the reads per million (RPM)-normalized piRNA abundance from each pachytene piRNA gene in CAST/EiJ compared to F1 hybrid males produced from reciprocal crosses of CAST/EiJ and C57BL/6J. Each dot represents the average of three biological replicates for CAST/EiJ and six biological replicates for F1 hybrid males. **(b)** Integrative Genomics Viewer (IGV) view illustrates RPM-normalized piRNA abundance from *pi17-CAST* locus in the testes from C57BL/6J, CAST/EiJ, and F1 hybrid males. Other replicates are reported in Extended Data Fig. 6b. **(c)** Histogram shows the RPM-normalized abundance of piRNAs that map to each CAST/EiJ- or C57BL/6J-specific single nucleotide polymorphism (SNP) in the testes from C57BL/6J, CAST/EiJ, and F1 hybrid males. Eutherian *2-qE1-35981* locus served as control (top). **(d)** Boxplots show CAST/EiJ allele frequency at *2-qE1-35981* (top) and *pi17-CAST* (bottom) loci in the testes from C57BL/6J, CAST/EiJ, and F1 hybrid males. CAST/EiJ allele frequency is calculated by computing the fraction of piRNA reads retaining CAST/EiJ-specific SNPs. Vertical lines represent median. Whiskers show 95% confidence intervals. Interquartile range (IQR) is represented by boxplots. Significance is measured by two-sided Wilcoxon matched-pairs signed rank sum test.

### Broad histone butyrylation over the *pi17-CAST* locus is maintained on CAST/EiJ allele in hybrid males

Specific to pachytene piRNA genes, the antagonism between skipped splicing and pronounced transcriptional elongation is partly explained by broad histone acylation that exceeds promoter-proximal region downstream and spreads over the gene body^50^. As in C57BL/6J, broad histone acylation the over gene body is a conserved feature of pachytene piRNA genes across house mice subspecies. Our CUT&RUN for histone H4 lysine 5 butyrylation (H4K5bu) from P/D spermatocytes of PWK/PhJ and CAST/EiJ mice revealed that the H4K5bu signal accumulated broadly over the gene bodies of pachytene piRNA genes (Extended Data Fig. 7a, top), whereas H4K5bu signal was confined to the promoters of other genes that are not making piRNAs and are regulated by A-MYB/TCFL5 regulatory axis^36^ (Extended Data Fig. 7a, bottom).

In CAST/EiJ, H4K5bu signal spread broadly over the gene body of *pi17-CAST* but not over that of *Lemd2* (Extended Data Fig. 7b). In contrast, *pi17-CAST* locus lacked H4K5bu signal in C57BL/6J (Extended Data Fig. 7b), consistent with its inactive transcription in C57BL/6J (Fig. 3b). Notably, P/D spermatocytes of F1 hybrid males accumulated a broad H4K5bu signal over *pi17-CAST* locus albeit at a lower density compared to CAST/EiJ (Extended Data Fig. 7b). This promoted us to investigate whether H4K5bu signal is accumulated on the CAST/EiJ allele in F1 hybrid males. We identified confident nucleotide variants between C57BL/6J and CAST/EiJ using H4K5bu CUT&RUN reads and measured the abundance of reads mapping to subspecies-specific SNPs in CAST/EiJ, F1 hybrids and C57BL/6J mice. *pi2* and *pi7* served as control. In F1 hybrids, the majority of H4K5bu CUT&RUN reads with subspecies-specific SNPs corresponded to CAST/EiJ allele of the *pi17-CAST* locus. On the contrary, for *pi2* or *pi7* loci, reads with subspecies-specific SNPs were equally distributed between C57BL/6J and CAST/EiJ (Extended Data Figure 7c). Consistent with this observation, CAST/EiJ allele frequency at *pi17-CAST* locus was nearly twice that at *pi2* or *pi7* genes in the F1 hybrids (Extended Data Fig. 7d; *pi17-CAST*, median = 0.71-1; mean = 0.68-0.83; *pi2,* median = 0.5; mean = 0.49-0.51; *pi7*, median = 0.4-1; mean = 0.38-0.56; two-sided wilcoxon matched-pairs signed rank sum test). Together with piRNA-based nucleotide variant analysis, we thus conclude that CAST/EiJ-specific nucleotide variants within *pi17-CAST* locus is essential for piRNA production.

### Hybrid males reveal reduced fertility features

The “Dobzhansky–Muller model” postulates that when two geographically isolated species inter-breed, negative epistatic interaction between different alleles of at least two loci from different species builds postzygotic reproductive isolation, ultimately resulting in hybrid sterility or breakdown^51,52^. Many piRNA–target pairs appear to be species-specific (Fig. 2e) implying that highly labile pachytene piRNAs and their target transcripts may represent a negative epistatic interaction in hybrid mice. To test this, we assessed fertility features of hybrid males from inter-breeding of C57BL/6J, PWK/PhJ, and CAST/EiJ (Extended Data Fig. 8) and investigated the piRNA–target pairs in hybrid males compared with parents (Fig. 5).

**Figure 5.**
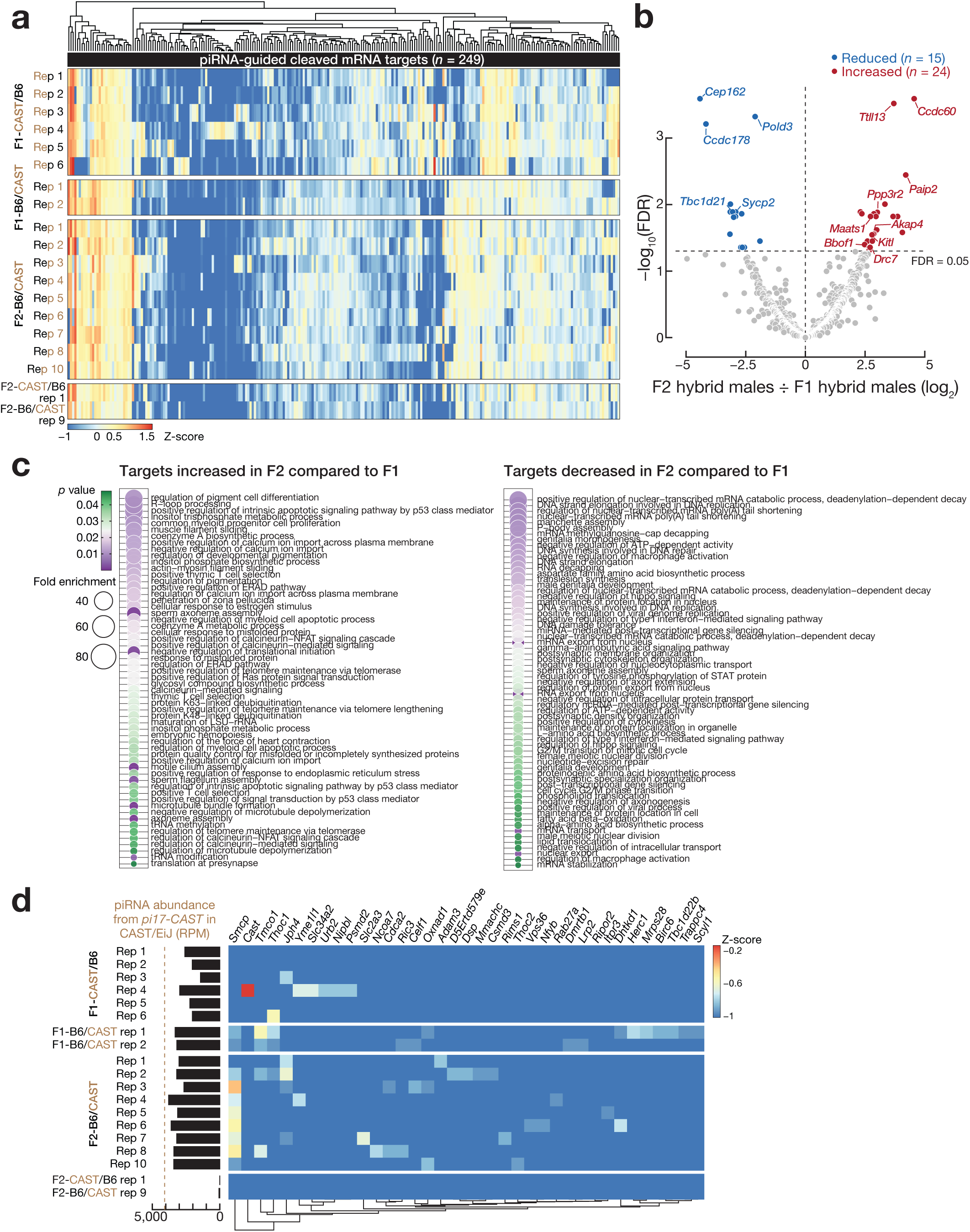
Distinct pachytene piRNA-mRNA pairs are gained or lost in F2 hybrid males compared to F1 hybrid males. **(a)** Heatmap representation of unsupervised hierarchical clustering of samples based on the abundance of piRNA-guided cleaved mRNA fragments (*n* = 249). **(b)** Volcano plot shows piRNA-guided cleaved mRNA fragments whose abundance is increased, decreased, or unchanged in F2 hybrid males (*n* = 11) compared to F1 hybrid males (*n* = 8). Blue denotes cleaved mRNA fragments whose abundance is reduced, while red represents those that are increased in F2 hybrid males. Differentially cleaved mRNAs (FDR < 0.05) were identified using DESeq2^67^. Benjamini-Hochberg correction was used to compute the false discovery rate. **(c)** Gene ontology (GO) analyses for piRNA-guided cleaved mRNA fragments whose abundance is increased (left) and decreased (right) in F2 hybrid males compared to F1 hybrid males. The gene categories are sorted by fold enrichment (*p* values <0.05 and FE > 20). See Supplementary Table 8 for GO terms with >2 fold enrichment. **(d)** Heatmap shows the change in the abundance of 37 mRNA fragments cleaved by piRNAs from *pi17-CAST* locus across F1 and F2 hybrid males. Bar plot represents the reads per million (RPM)-normalized abundance of piRNAs from *pi17-CAST* locus in each sample.

As in previous studies^19–22,53^, F1 hybrid males from the C57BL/6J ♂ × PWK/PhJ ♀ crosses (Henceforth, F1-B6/PWK) displayed sterility (Extended Data Fig. 8). F1-B6/PWK males failed to sire any litters, whereas F1 males from the opposite cross, PWK/PhJ ♂ × C57BL/6J ♀ (Henceforth, F1-PWK/B6), were fertile and sired F2 litters (Extended Data Fig. 8a–c). Consistently, testis mass and sperm quantity from the caudal epididymis were markedly reduced in F1-B6/PWK compared with either parental strain (Extended Data Fig. 8b, c; Kruskal-Wallis test, pairwise Wilcoxon–Mann-Whitney test with Benjamini-Hochberg correction, adjusted *p* values; testis mass, median F1-B6/PWK = 0.11 ± 0.02 g, median C57BL/6J = 0.21 ± 0.01, median PWK/PhJ = 0.19 ± 0.01; sperm quantity, median F1-B6/PWK = 0.14 ± 0.3 million sperm per ml, median C57BL/6J = 2.2 ± 0.9, median PWK/PhJ = 1.5 ± 0.9).

F1 hybrid males from the reciprocal crossing C57BL/6J and CAST/EiJ were fertile and sired F2 litters (Extended Data Fig. 8b, c). Indeed, regardless of the direction of crossing, both sperm velocity and the fraction of fast sperms with >100 µm × s^-1^ in F1 hybrid males appeared to be higher than their parental inbred strains (Extended Data Fig. 8d–f). Notably, however, F2 hybrid males from the C57BL/6J ♂ × CAST/EiJ ♀ crosses (Henceforth, F2-B6/CAST) exhibited defective sperm features (Extended Data Fig. 8d–f). A similar trend was observed in F2 hybrid males from the opposite crosses (i.e., F2-CAST/B6). Sperm populations from F2-B6/CAST exhibited significantly reduced path and progressive velocities when compared with F1-B6/CAST (Extended Data Fig. 8e). Furthermore, the fraction of fast sperms with >100 µm × s^-1^ was markedly reduced in F2-B6/CAST compared to F1-B6/CAST (Extended Data Fig. 8f). Together, F2 hybrid males from the reciprocal crossing C57BL/6J and CAST/EiJ exhibited defective fertility features, consistent with the studies showing that sperm fitness decreases over hybrid generations, i.e., hybrid breakdown^54,55^.

### Distinct pachytene piRNA-mRNA pairs are gained or lost in F2 hybrid males when compared to F1 hybrid males

To examine whether pachytene piRNAs have distinct sets of targets in F2 hybrid males with reduced fertility features compared to F1 hybrid males, we sequenced 5’-monophosphorylated long RNAs and piRNAs of F1-B6/CAST, F1-CAST/B6, F2-CAST/B6, and F2- B6/CAST males. Cleavage products were required to have g2 to g15 perfect complementarity to piRNAs of hybrid males (Methods). Here, our analysis defined 249 putative mRNA targets (Fig. 5a). Unsupervised hierarchical clustering revealed distinct patterns of the cleaved mRNA-fragments abundance across hybrid males (Fig. 5a).

The direction of reciprocal crossing between C57BL/6J and PWK/PhJ had different outcome: F1-B6/PWK males were sterile, while F1-PWK/B6 males were fertile (Extended Data Fig. 8). By contrast, directionality had no impact on the fertility features of F1 hybrid males produced from the crossing between C57BL/6J and CAST/EiJ.

Indeed, both F1-B6/CAST and F1-CAST/B6 males were fertile and sired F2 litters (Extended Data Fig. 8). However, regardless of the directionality, F2 hybrid males exhibited reduced fertility features (Extended Data Fig. 8). We thus performed DESeq2 analysis to define the differentially cleaved mRNA-fragments between F1 and F2 hybrid males (Fig. 5b). The abundance of 39 piRNA-guided cleaved mRNA-fragments were significantly different in F2 hybrid males compared with F1 hybrid males (Fig. 5b; Supplementary Table 7; F2 hybrid males ÷ F1 hybrid males ≤ 0.5 or ≥ 2; FDR < 0.05).

Of the 39 differentially cleaved mRNA-fragments, the abundance of 24 was increased, whereas 15 were lost/decreased in F2 hybrid males (Fig. 5b). Notably, differentially cleaved mRNA-fragments were enriched for gene categories that included sperm axoneme and flagellum assembly, manchette assembly, and motile cilium assembly (Fig. 5c; Supplementary Table 8; Gene Ontology analysis; GO; ≥2-fold change; *p* value < 0.05).

Finally, we examined whether piRNAs from *pi17-CAST* locus have distinct targets in F2 hybrid males when compared to F1 hybrid males. Interestingly, 37 differentially cleaved mRNA fragments included *Smcp* mRNA encoding a protein required for structural organization of the sperm mitochondrial sheath^56^ (Fig. 5d). *Smcp* mRNA was cleaved in F2 hybrid males that produce *pi17-CAST* piRNAs, whereas F1 hybrid males showed almost undetectable *Smcp* cleavage (Fig. 5d). Indeed, F2 hybrid males that do not produce *pi17-CAST* piRNAs equally failed to cleave *Smcp*, suggesting that *pi17-CAST* piRNA dependent cleavage of *Smcp* mRNA is gained in F2 hybrid males (Fig. 5d). Together, our findings reveal that over the course of two generations, hybrid males either gained new piRNA-mRNA pairs or lost existing piRNA-mRNA pairs.

## Discussion

Pachytene piRNA genes are required for mammalian male reproduction and evolve rapidly^4,5,10,12,18^. Whether high evolutionary turnover of pachytene piRNAs is beneficial their function has remained elusive, partly due to the lack of an experimental mammalian system that enables direct investigation of the evolutionary role of pachytene piRNAs. Combining multiple genomic approaches, we report precise genomic coordinates of pachytene piRNA genes in the genomes of geographically isolated two mouse subspecies, PWK/PhJ (*Mus musculus musculus*) and CAST/EiJ (*Mus musculus castaneus*).

The transcriptional machinery driving the expression of pachytene piRNAs is deeply conserved. In both PWK/PhJ and CAST/EiJ, promoters of >90% of pachytene piRNA genes are under the control of A-MYB/TCFL5 regulatory axis (Extended Data Fig. 3), mirroring C57BL/6J (*Mus musculus domesticus*)^11,17^. We recently reported that A-MYB-dependent eutherian pachytene piRNA genes, but not younger loci, exhibit broad histone acylation over their gene bodies^14,17^. Moreover, deeply conserved loci produce more piRNAs than younger loci^12,43^, and tend to have bidirectional promoters^14^. Together, these findings suggest that, over long evolutionary timescale, both the acquisition of strong A-MYB occupancy at the promoter-proximal region and accumulation of histone acylation over the gene body establish a locus as a highly prolific piRNA source. However, the CAST/EiJ-specific *pi17-CAST* gene defies this notion entirely. We show that the abundance of its piRNAs surpasses the median abundance of conserved loci (Fig. 3), and that *pi17-CAST* is transcribed divergently from an A-MYB-dependent bidirectional promoter that generates *Lemd2* mRNA from the opposite arm. Intriguingly, in F1 hybrid males, piRNAs are produced almost exclusively from the CAST/EiJ allele (Fig. 4; Extended Data Fig. 8). Bidirectional promoters are abundant throughout mammalian genome^57^. In male germ cells, the A-MYB/TCFL5 circuit regulates many bidirectional promoters with no piRNA production at both opposite arms^17,41^, suggesting that promoter-proximal regions alone are insufficient to turn a locus into a piRNA source. Could there then be undefined genetic elements beyond promoter-proximal regions specific to pachytene piRNA genes, perhaps within gene body? *pi17-CAST* gene therefore provides an excellent opportunity for future studies elucidating cis-acting genetic elements that are specific to pachytene piRNA genes.

Genomic locations of all pacyhtene piRNA genes are conserved across mice subspecies, yet 16 loci produce pachytene piRNAs subspecies specifically underscoring how labile pachytene piRNA genes are during a short evolutionary timescale (∼0.5 million years ^23–25^). Exons of such species-specific loci diverge faster than that of loci found in all three subspecies (Fig. 2), suggesting that newly acquired piRNA sequences presumably regulate different targets in emerging species to regulate the sperm fitness within that species. In fact, male mice expressing a human pachytene piRNA gene become infertile due to human piRNAs apparently dysregulating mRNAs that are not supposed to be silenced in mouse^58^. We report that CAST/EiJ innovated *pi17-CAST* locus producing abundant pachytene piRNAs. Exciting future experiments, such as assessing the fertility features and piRNA-target pairs of C57BL/6J males ectopically expressing *pi17-CAST* locus, will enable us to directly investigate the role of a newly emerged pachytene piRNA gene in mammalian reproductive barrier.

The “Dobzhansky–Muller model” predicts that different alleles of two epistatically interacting loci become incompatible in the hybrids of related species^51,52^. To date, in mammals, only one speciation gene, *Prdm9*, and its incompatibility with X-linked loci, including *Hstx2*, is mapped. Such incompatible loci were characterized in F1 hybrid males produced from the crossing of C57BL/6J males and PWK/PhJ females^19–22^.

However, when fertile F1 females are backcrossed to either parental males or are intercrossed with partially fertile F1 males can result in F2 hybrid males that reveal compromised fertility features^20^. This suggests that reproductive barrier is multigenic involving recessive epistatic incompatibilities. Such recessive incompatible loci thus remain to be identified. piRNA-guided mRNA cleaved fragments specifically identified in F2 hybrid males with reduced sperm fitness presumably exemplifies recessive epistatic incompatibilities. That is, distinct piRNA sequence expressed from one of the parental haplotypes may potentially direct the cleavage of mRNA variant that is produced from the opposite parental haplotype. Indeed, we found that *Smcp* mRNA was only cleaved in F2 hybrid males producing *pi17-CAST* piRNAs but not in F1 hybrid males despite the expression of *pi17-CAST* locus (Fig. 5d). Furthermore, *Smcp* mRNA was not cleaved by *pi17-CAST* piRNAs in CAST/EiJ males either (Fig. 2; Supplementary Table 6). These observations suggest that the emergence or loss of piRNA–target pairs may reflect diverging sequences more likely in piRNAs than in mRNAs because pachytene piRNA sequences evolve faster than other genomic elements (Fig. 2; Extended Data Fig. 4) However, we cannot exclude the possibility that F2 hybrid males express *Smcp* mRNA with C57BL/6J-specific nucleotide variants that become a target for *pi17-CAST* piRNAs. Together, our findings provide the evidence that rapidly diverging pachytene piRNAs can gain or lose targets in the hybrid males of closely related mammalian species.

## Methods

### Mice

C57BL/6J (RRID: IMSR_JAX:000664), PWK/PhJ (RRID: IMSR_JAX:003715), and CAST/EiJ (RRID: IMSR_JAX:000928) mice were included in this study. Mice were maintained and used according to the guidelines of the Regional Animal Experimentation Ethics Committee of Swedish Board of Agriculture with the animal ethical permit numbers 8246-2021 and 5.8.18-3736/2026.

### Mouse male fertility analysis

#### Sperm count

To quantify number of sperms, caudal epididymides were collected from 2–12-month-old mice and placed in Ham’s F-10 medium (Gibco; 22390) supplemented with 10% bovine serum albumin (Sigma-Aldrich; A9418). Several small incisions were made in the epididymides using scissors in order to release sperm into the medium that was incubated at 37 °C in 5% CO₂ for 20 min. The sperm suspension was diluted in 2.5–5% (v/v) formaldehyde, and sperm cells were counted under bright-field microscopy.

#### Motility assay

For sperm motility analysis, a drop of sperm suspension was loaded into a sperm-counting chamber (ISAS; D4C20L) and analyzed using a computer-assisted sperm analysis (CASA) system (UB 200i Series Microscope and C13-ON camera, PROiSER R+D). The following settings were used: species, rodent; 60 acquired images at 60 frames/second (s); optics, phase contrast (Ph−); chamber type, ISAS-D4C20; magnification, ×10; particle area, 6–70 µm²; slow cells, <10 µm/s; medium cells, 15–45 µm/s; rapid cells, >45 µm/s; progressivity threshold, 50% STR; connectivity, 8; and a minimum of 10 images to calculate ALH. For sperm motility quantification, sperms whose movement was captured with ≥30 frames were used for analysis.

### Isolation of mouse germ cells by Fluorescence-Activated Cell Sorting

Male germ cell sorting was performed as previously described using BD FACSMelody Cell Sorter (BD Biosciences)^36^. In brief, seminiferous tubules were squeezed into 6 ml 1× Gey′s Balanced Salt Solution (GBSS, Sigma; G9779) containing 0.4 mg / ml collagenase type IV (Worthington; LS004188) and incubated at 33°C for 15 min at 600 rpm. Thereafter, the tubules were washed twice with 6 ml GBSS, which was followed by incubation in 1× GBSS containing 0.5 mg/ml trypsin (Gibco; 27250-018) and 1 µg/ml DNase I (Roche; 10104159001) at 33°C for 15 min at 600 rpm. The tubules were then gently homogenized at 4°C on ice using a Pasteur pipette. We added 400 μl FBS to inactivate trypsin. The cells were then passed through a pre-wetted 70 μm cell strainer (Corning; CLS431751). Afterwards, cells were pelleted in swing bucket centrifuge at 500 × g for 10 min at 4°C. The cells were resuspended in 1× GBSS containing 5% (v/v) FBS, 1 µg/ml DNase I, and 5 μg/ml Hoechst 33342 (ThermoFisher; 62249) and incubated at 33°C for 45 min at 150 rpm. Propidium iodide (PI; 0.2 μg/ml, f.c.; ThermoFisher; P3566) was added to the cell suspension and the cells were finally filtered through a pre-wetted 40 µm cell strainer (Corning; CLS431750). Four-way cell sorting was performed to obtain the populations of spermatogonia (SpG), leptotene and zygotene (L/Z), pachytene and diplotene (P/D), and round spermatids (RS) on BD FACSMelody Cell Sorter (BD Biosciences) using the gates as described previously^36^.

### Long RNA library construction and analysis

Total RNA was extracted using mirVana miRNA RNA isolation kit (ThermoFisher; AM1561). Library construction was performed as previously described^12,36,59^. In brief, 2 μg total RNA was mixed with 2 μl antisense DNA oligonucleotides (a pool of 186 different oligos that are complementary to ribosomal RNA [rRNA] sequences; 0.5 μM each^60,61^) in 10 μl with rRNA oligo hybridization buffer (100 mM Tris-HCl [pH 7.4] and 200 mM NaCl). The mixture was incubated at 95°C for 5 min and cooled gradually to 25°C at a rate of –0.1°C per second, and held at 25°C for 5 min. Afterwards, the DNA:RNA hybrids were degraded by incubating the mixture with 10 U Thermostable RNase H (Epicenter; E0038-5D1) in 50 mM Tris-HCl (pH 7.4), 100 mM NaCl, 20 mM MgCl_2_ at 45°C for 30 min. Subsequently, the samples were incubated with 4U Turbo DNase (ThermoFisher; AM2238) at 37°C for 20 min and purified using the RNA Clean & Concentrator kit (Zymo Research; R1015). End-repair, A-tailing, adaptor ligation, UDG treatment and library amplification were performed as previously described^12,36,59^. Libraries were sequenced as paired-end reads using NextSeq550, NextSeq2000 or NovaSeqX Plus system (Illumina).

We first removed all RNA-seq reads that mapped to ribosomal RNAs using (v2.5.5^62^) with default parameters. The remaining reads were mapped to the reference mouse genome (mm10) using STAR (v2.2.1^63^). Mapped reads were then assigned to protein-coding genes, non-coding RNAs, and piRNA genes using BEDTools (v2.27.1^64^) and normalized to reads per kilobase of transcript per million (RPKM).

### piRNA library construction and analysis

Total RNA was isolated using the MirVana RNA isolation kit (ThermoFisher; AM1561). piRNA libraries were constructed as previously described^12^. Briefly, 18–36 nt small RNAs were size selected by separating 2 μg total RNA on 15% denaturing urea polyacrylamide gel electrophoresis. To exclude other small RNAs from the library, RNA was oxidized with 30 mM (w/v) borax (ThermoFisher; A16176), 30 mM (w/v) boric acid (ThermoFisher; 012680), (pH 8.6) and 25 mM (w/v) sodium periodate (ThermoFisher; 198381000) at 25°C for 30 min before 3’ adaptor ligation. The oxidized small RNAs were ligated to AppBA3-UMI 3’ adaptor (5′-rAppNNNGTCNNNTAGNNNTGGAATTCTCGGGTGCCAAGG/ddC/-3) in 50 mM (w/v) Tris -HCL (pH 7.5), 10 mM (w/v) MgCl2, 10 mM (w/v) DTT, 25% PEG8000, and 600 U T4 Rnl2tr K227G (NEB; M0351S). The 3’ ligation reaction was incubated at 16°C for 16 h. Adaptor ligated small RNAs (63-65 nt) were then purified on 15 % denaturing urea polyacrylamide gel electrophoresis. The samples were then incubated at 90°C for 1 min then at 65°C for 5 min before BRTP (5’-CCTTGGCACCCGAGAATTCCA-3’) primer was added. The samples were afterwards incubated at 65°C for 5 min. Thereafter, BA5-UMI 5’ adaptor (12.5 μM 5’ - GUUCAGAGUUCUACAGUCCGACGAUCNNNCGANNNUACNNN-3’, 12.5 μM 5’ - GUUCAGAGUUCUACAGUCCGACGAUCNNNAUCNNNAGUNNN-3’) was ligated to small RNAs in ligation buffer (50 mM (w/v) Tris -HCL [pH 7.5], 10 mM (w/v) MgCl2, 10 mM (w/v) DTT, 1 mM ATP) containing 10 U T4 RNA ligase (Ambion; AM2141) at 25°C for 2 h. Reverse transcription reaction was performed in 1X AMV RT buffer (NEB; B0277A) containing 1.5 mM dNTP mix and 20 U AMV Reverse Transcriptase (NEB; M0227) at 42°C for 1h, and 90°C for 5 min followed by ice incubation. Small RNA libraries were then amplified in 1× AccuPrime *pfx* reaction mix (Invitrogen; 12344-024), containing 0.5 μM BPCRP1 primer (5′-AATGATACGGCGACCACCGAGATCTACACGTTCAGAGTTCTACAGTCCGA-3′), 0.5 μM PCRIdx (5′- CAAGCAGAAGACGGCATACGAGATNNNNNNGTGACTGGAGTTCCTTGGCACCCGA GAATTCCA-3′), and 1.25 U AccuPrime *pfx* DNA polymerase (Invitrogen; 12344-024). The following PCR condition was applied to generate small RNA libraries: 95°C for 2 min; 15 cycles of thermocycling at 95°C for 15 sec, 65°C for 30 sec; and 68°C for 15 sec. 160–180 bp small RNA libraries were finally purified on 2% (w/v) agarose gel or 8% native polyacrylamide gel electrophoresis. Small RNA libraries were sequenced single-end reads on NextSeq550 or NextSeq2000 system (Illumina).

We first removed the PCR duplicates using Umitools^65^ and adapter sequences from all small RNA-seq reads using Cutadapt (v5.2^66^) with default parameters. Reads shorter than 18-nt or longer than 32-nt were filtered out prior to genome alignment.

Subsequently, reads mapping to ribosomal RNAs, miRNAs, snoRNAs, snRNAs, and tRNAs were excluded. The remaining small RNA-seq reads were independently mapped to the reference mouse genome (mm10) with parameters `-v 1 -a -best - strata`. Lastly, we quantified piRNA abundance normalized by sequencing depth that was defined as the total number of uniquely genome mapping reads after the removal of rRNAs, miRNAs, snoRNAs, snRNAs, and tRNAs.

### Degradome sequencing and analysis

Libraries for 5’ monophosphorylated 3’-end cleavage products were prepared as previously described^10^. In brief, total RNA was isolated from frozen testis using MirVana kit (ThermoFisher; AM1561). rRNA sequences were depleted from 4 μg total RNA described previously (Long RNA library construction). 5’ adaptor, whose 3’-end is with hydroxyl, was ligated in T4 RNA ligase buffer (0.5 mM (w/v) Tris [pH 7.8], 0.1 mM (w/v) MgCl2, 0.1 mM (w/v) DTT, 0.01 mM (w/v) ATP) containing 1.25 μM BA5-UMI adaptors and 10 U T4 RNA ligase (Ambion; AM2141) at 25°C for 2 h. After 5’ adaptor ligation, RNA was purified using RNA Clean & Concentrator (Zymo Research; R1015).

Afterwards, complementarity DNA was generated using random Deg-RT primer (5’–GCACCCGAGAATTCCANNNNNNNN–3’), which retains the sequence of 3’ adaptor. Reverse transcription was performed using 200 U SuperScript III (ThermoFisher; 18080093) with the following program: 25°C for 5 min; 50°C for 1 h; 70°C for 15 min. Next, libraries were amplified in 1× AccuPrime buffer (ThermoFisher; 12344024), 0.2 μM Deg-PCR-1L (5’–CTACACGTTCAGAGTTCTACAGTCCGA–3’), 0.2 μM Deg-PCR-1R (5’–GCCTTGGCACCCGAGAATTCCA–3’), and 1.25 U AccuPrime pfx (ThermoFisher; 12344024) with the following PCR condition: 98°C for 30 s, two cycles of thermocycling at 95°C for 15 s, 59°C for 30 s, 68°C for 12 s; four cycles of thermocycling at 95°C for 10 s, 68°C for 22 s; 68°C for 3 min. 200–500 bp PCR product was then purified on 8% native polyacrylamide gel electrophoresis. Second PCR was performed in 1× AccuPrime buffer (ThermoFisher; 12344024), 10.2 μM BPCRP1 primer, 0.2 μM BPCRPIdX primer, 1.25 U AccuPrime pfx (ThermoFisher; 12344024) with the following PCR condition: 95°C for 30 s; two cycles of thermocycling at 95°C for 10 s, 65°C for 30 s, 68°C for 30 s; 10 cycles of thermocycling at 95°C for 10 s, 68°C for 30 s; final extension at 68°C for 3 min. 250–550 bp final libraries were purified on 8% native polyacrylamide gel electrophoresis. Libraries were sequenced paired-end on NextSeq550 or NextSeq2000 system (Illumina).

We first aligned degradome-seq reads to genome as previously described for RNA-seq. Specifically, we removed all degradome-seq reads that mapped to ribosomal RNAs using Bowtie2 (v2.5.4^62^) and mapped the remaining reads to the mouse reference genome (mm10) using STAR (v2.2.1^63^) with default parameters. Afterwards, read1 from the read pairs with MAPQ ≥ 10 and no soft-clipping were retained. The 5’-end of the read1 was used to define guide sequences, which were extended 10 base pairs upstream and 40 base pairs downstream. Those guide sequences, which perfectly matched to piRNA reads from the guide nucleotides g2 to g15 positions, were defined as piRNA-target pairs. The abundance of cleaved products (degradome reads) was normalized to reads per million filtered read1. Differentially cleaved mRNA-fragments between F2 hybrid males and F1 hybrid males were defined using DESeq2^67^.

### CUT&RUN sequencing and analysis

<u>C</u>leavage <u>U</u>nder <u>T</u>argets and <u>R</u>elease Using Nuclease (CUT&RUN) sequencing from FACS-purified P/D spermatocytes was performed according to the manufacturer’s protocol (Active Motif; 53180) as described previously^36^. 500,000 FACS-purified P/D cells were gently resuspended in 100 μl nuclear isolation buffer (20 mM [w/v] HEPES-KOH, pH 7.9, 10 mM [w/v] KCl, 0.5 mM [v/v] Spermidine, 0.1% [v/v] Triton X-100, 20% [v/v] glycerol, 1× protease inhibitors, 0.5 mM Spermidine) and incubated on ice for 10 min. Afterwards, cells were centrifuged at 600×g at 4°C for 3 min. Nuclei were then washed once and resuspended in 100 μl wash buffer to which 10 μl Concanavalin A-coated paramagnetic beads was added. To immobilize the nuclei on beads, we incubated the mixture at room temperature for 10 min. After incubation, supernatant was removed using magnet and 50 μl antibody buffer containing 2 μg anti-MYBL1 (Sigma; HPA008791), or anti-TCFL5 (Sigma; HPA055223), or 1 μl anti-H4K5bu (PTM bio; PTM-310), or 1 μl IgG (EpiCypher; 13-0042) antibodies was added to beads. Samples were incubated at 4°C overnight on a nutator. After washing the beads twice in 200 μl cell permeabilization buffer (CPB), 50 μl CPB containing 2.5 μl ChIC/CUT&RUN pAG-MNase was added to beads and incubated at room temperature for 10 min. After incubation, following two additional washes in 200 μl CPB, we added 1 μl 0.1 M CaCl_2_ to start tagmentation at 4°C for 2 hours on a nutator. The reaction was stopped by adding 40 μl STOP solution and incubated at 37°C for 10 min in a thermal cycler.

Samples were then placed on magnet and the supernatant containing the DNA fragments were moved to a new tube. After purifying the DNA using columns provided with the kit, libraries were prepared using NEBNext Ultra II DNA Library Prep Kit (NEB; E7645) according to the manufacturer’s protocol. CUT&RUN libraries were sequenced as paired-end reads on NextSeq550 or or NovaSeq X Plus (Illumina).

Adapters were trimmed from the quality filtered paired-end reads using Fastp (v0.24.0^68^). Afterwards, reads were aligned to mouse reference genome (mm10) using Bowtie2 (v2.5.4^62^) with the parameters that allows fragment sizes between 10 and 700 bp `-local -very-sensitive -no-unal-no-mixed -no-discordant -|10 - X700`. Alignments were converted to BAM format, sorted, and indexed using SAMtools (v1.8^69^). Only uniquely mapped reads (MAPQ ≥ 20) were retained for downstream analyses. PCR duplicates were removed using Picard MarkDuplicates with parameters `REMOVE_DUPLICATES=true VALIDATION_STRINGENCY=LENIENT`. Resulting BAM files were indexed. Alignment statistics were obtained with `samtools flagstat`. Genome-wide coverage tracks were generated from duplicate-filtered BAM files using bamCoverage from the deepTools package (v3.5.5^70^). Reads per million (RPM)-normalized BigWig files were generated using a bin size of 25 bp with read extension enabled. For H4K5bu (Extended Data Fig. 7a) CUT&RUN, corresponding subspecies-matched IgG controls were subtracted using `bigwigCompare -operation subtract`. Following IgG subtraction, biological replicates were merged by averaging the signal across two replicates using the UCSC `bigWigMerge` utility (v2). The resulting bedGraph files were converted back to BigWig format using `bedGraphToBigWig` (v2.8).

A-MYB or TCFL5 peaks were identified using MACS3 (v3.0.2^71^; FDR < 0.05). CUT&RUN for IgG antibody was used as the background control to call significant A-MYB or TCFL5 peaks. Genes with A-MYB or TCFL5 peak within ±2 kb of their transcription start sites were considered as A-MYB- or TCFL5-bound genes.

### qPRO sequencing (qPRO-seq) and analysis

#### Nuclei isolation

1,000,000 P/D spermatocytes were pelleted at 400 × g at 4°C for 10 minutes. After removing the supernatant carefully, the cell pellet was resuspended in 650 μl swelling buffer (10 mM (w/v) Tris-HCl pH 7.5, 2 mM (w/v) MgCl2, 3 mM (w/v) CaCl2), to which we then added additional 6 ml swelling buffer slowly. Cells were then incubated on ice for 5 min and pelleted at 400 × g at 4°C for 10 min. The cell pellet was resuspended in 335 μl lysis buffer 1 (9 mM (w/v) Tris-HCl [pH 7.5], 1.8 mM (w/v) MgCl2, 2.7 mM (w/v) CaCl2, 10% (v/v) glycerol, 1× protease inhibitor cocktail (PIC), 0.4 U/μl RNAsIn (Promega; N2615). Thereafter, 335 μl lysis buffer 2 (9 mM (w/v) Tris-HCl [pH 7.5], 1.8 mM (w/v) MgCl2, 2.7 mM (w/v) CaCl2, 10% (v/v) glycerol, 5% (v/v) IGEPAL CA-360, 1× PIC, 0.4 U/μl RNAsIn [Promega; N2615]) was added dropwise to the mix and incubated on ice for 5 min. After incubation, we slowly added 6 ml lysis buffer 3 (9 mM (w/v) Tris-HCl pH 7.5, 1.8 mM (w/v) MgCl2, 2.7 mM (w/v) CaCl2, 10% (v/v) glycerol, 0.005% (v/v) Igepal CA-360, 1× PIC, 0.4 U/μl RNAsIn [Promega; N2615]) to the nuclei that were then were pelleted at 600 × g at 4°C for 5 min. The supernatant was discarded and the nuclei were resuspended in 668 μl lysis buffer 3. Afterwards, we slowly added 6 ml lysis buffer 3 dropwise to the nuclei and carefully mixed. The nuclei were centrifuged at 500 × g at 4°C for 5 min. The supernatant was removed and the nuclei were resuspended in 668 μl freezing buffer (50 mM (w/v) Tris-HCl pH 8, 5 mM (w/v) MgCl2, 0.1 mM (w/v) EDTA pH 8, 40% (v/v) glycerol, 1× PIC, 0.4 U/μl RNAsIn [Promega; N2615]). The nuclei were centrifuged at 900 × g at 4°C for 6 min. The supernatant was discarded and the nuclei were finally resuspended in 50 μl freezing buffer.

#### Run-on reaction and library construction

100 μl reaction mixture containing 50 μl nuclei in freezing buffer and 50 μl ROMM (5 mM (w/v) Tris-HCl pH 8.5, 2.5 mM (w/v) MgCl2, 0.5 mM DTT, 150 mM (w/v) KCl, 250 μM Biotin-11-UTP/CTP, 250 μM ATP/GTP, 0.5% (w/v) sarcosyl) was incubated at 37 °C at 750 RPM for 5 min. To stop the reaction, 350 μl RL buffer (NORGEN RNA extraction kit; 37500) was added. We then added 240 μl 100% ethanol to mixture and passed it through an extraction column at 3,500 x g for 1 min. The filter was washed twice with 400 μl wash solution A via centrifugation at 14,000 x g for 1 min. The column was dried by additional spinning at 14,000 x g for 2 min. RNA was then eluted twice in 50 μl water and denatured at 65°C for 30 s followed by cooling on ice. Thereafter, RNA was fragmented in 200 mM (w/v) NaOH on ice for 10 min. The reaction was stopped by adding 125 μl 1 M (w/v) Tris-Cl pH 6.8. We then cleaned RNA by ethanol precipitation. The precipitated RNA pellet was resuspended in 6 μl water. We added 1 μl 5 μM VRA3 adaptor to RNA and denatured the RNA at 65°C for 30 s that was followed by snap cooling on ice. 3’ adaptor ligation was performed in 1× T4 RNA ligase buffer (NEB; B0216S), 1 mM ATP, 20 U SUPERase-In (ThermoFisher; AM2696), 15% PEG8000, 20 U T4 RNA Ligase 1 [NEB; M0204S]) at 25°C for 1 hour. After ligation, ligated RNA was mixed with binding buffer (7.3 mM (w/v) Tris-HCl pH 7.4, 220 mM (w/v) NaCl, 0.073 % (v/v) Triton X-100, 0.73 mM (w/v) EDTA, with 0.004 U SUPERase-In). We then added 25 μl Streptavidin C1 beads (ThermoFisher; 65001) directly to the samples and incubated on rotor at room temperature for 25 min. The samples were washed once with 500 μl high salt buffer (50 mM (w/v) Tris-HCl pH 7.4, 2M (w/v) NaCl, 0.5% (v/v) Triton X-100,1 mM (w/v) EDTA, with 0.004 U SUPERase-In), and once with 500 μl low salt buffer (5 mM (w/v) Tris-HCl pH 7.4, 0.1% (v/v) Triton X-100, 1 mM (w/v) EDTA, with 0.004 U SUPERase-In). After washing, samples were resuspended in 19 μl PNK mix (1× PNK buffer [NEB; B0201S], 1 mM ATP, 10 U T4 PNK [NEB; M0201S], 20 U SUPERase-IN) and incubated at 25°C for 30 min. 5’ decapping was performed by adding 19 μl cap mix (1× Cap-Clip buffer (CELLSCRIPT; C-CC15011H), 5 U Cap-Clip [CELLSCRIPT; C-CC15011H], 20 U SUPERase-In) and incubated at 37°C for 1 h. After incubation, the samples were resuspended in 7 μl adapter mix (1.4 μM REV5) and denatured at 65°C for 30 s that was followed by cooling on ice. Thereafter, 12 μl ligation mix (1× T4 RNA ligase buffer [NEB; B0216S],1 mM ATP, 20 U SUPERase-In [ThermoFisher; AM2696],15% PEG8000, 10 U T4 RNA Ligase 1 [NEB; M0204S]) was added to samples and incubated at 25°C for 1 h. After ligation, RNA was washed once with 500 μl high salt buffer and once with 500 μl low salt buffer. Afterwards, we performed RNA extraction using TRIzol reagent (ThermoFisher; 10296028). The ethanol precipitated RNA pellet was resuspended in 13.5 μl RT resuspension mix (2.5 μM RP1 primer, 0.5 mM dNTP mix) and denatured at 65°C for 5 min, followed by cooling on ice. Complementarity DNA was generated in 6.5 μl RT master mix (1× RT buffer [ThermoFisher; EP0752], 5 mM (w/v) DTT, 10 U SUPERase-In, 200 U Maxima H Minus reverse transcriptase [ThermoFisher; EP0752]) with the following thermocycler program: 50°C for 30 min; 65°C for 15 min; 85°C for 5 min. Full scale amplification was performed immediately after revers transcription (RT). qPRO-seq libraries were then amplified in PCR master mix (1× Q5 buffer [NEB; B9027S], 1× Q5 enhancer [NEB; B9028S], 0.1 μM RP1, 0.25 mM dNTP mix, 2 U Q5 polymerase [NEB; M0491S]) containing 0.25 μM index RPI primers. The following PCR condition was applied to generate qPRO-seq libraries: 95°C for 2 min; 5 cycles of thermocycling at 95°C for 30 s, 56°C for 30 s, 72°C for 30 s; 14 cycles of thermocycling at 95°C for 30 s, 65°C for 30 s, 72°C for 30 s. qPRO-seq libraries were sequenced as paired-end reads on NextSeq550 (Illumina).

qPRO-seq analysis was performed as previously described^36^. Briefly, adapter sequences were trimmed using fastp (v0.23^68^). Subsequently, reads were aligned to mouse reference genome (mm10) using Bowtie2 (v2.5.4^62^) with the parameters `—local —very-sensitive —no-unal —no-mixed —no-discordant -|10 -X700`. SAM files were converted to BAM format using SAMtools (v1.20^69^). The resulting BAM files were sorted and indexed, and uniquely mapped reads were retained by filtering alignments with a mapping quality (MAPQ) ≥ 20 using Samtools view `-F 0x904 -q 20`. Final indexed BAM files were then converted into reads per million (RPM)-normalized bigWig files using deepTools (v3.5.5^70^).

### Poly(A) Site sequencing (PAS-seq) and analysis

Poly(A) Site Sequencing (PAS-seq) was performed as previously described^12^. Briefly, total RNA was isolated from 500,000 P/D spermatocytes using MirVana RNA isolation kit (ThermoFisher; AM1561). We extracted poly(A)+ RNA from 1 μg total RNA using Dynabeads™ mRNA Purification Kit (Invitrogen; 61006). We next fragmented poly(A)+ RNA samples in 5 μl RNA fragmentation reagent (Invitrogen; AM8740) at 70°C for 5 min. To stop the reaction, 5.5 μl Stop Solution (Invitrogen; #AM8740) was added and placed on ice for 2 min. Fragmented RNA samples were ethanol precipitated and the precipitated RNA pellet was dissolved in water. Thereafter, First-strand buffer (Invitrogen; 18080093) containing 1 mM dNTP mix and 1 μM PAS-seq2 oligo(dT) primer (5’-GTGACTGGAGTTCAGACGTGTGCTCTTCCGATCTTTTTTTTTTTTTTTTTTTTV-3’) was added to the RNA samples and heated at 72°C for 3 min, followed by cooling on ice. We then added 5 mM DTT, 1 M Betaine (Sigma-Aldrich; B0300-1VL), 6 mM MgCl_2_, and 1 μM template switch oligo containing locked nucleic acid (5’-CTACACGACGCTCTTCCGATCTCATrGrG+G-3’) and 200 U SuperScript III (Invitrogen; 18080093) to perform reverse transcription in thermocycler with the following program: 42°C for 90 min, nine cycles of thermocycling at 50°C for 2 min, 42°C for 2 min; 72°C for 15min. The cDNA was finally amplified in HF buffer (NEB; B0518S), 0.2 mM dNTP mix, 0.2 μM TruSeq universal adapter, 0.2 μM TruSeq indexed adapter, and 1 U Phusion polymerase (NEB; M0530S). The following PCR condition was applied to generate PAS-seq libraries: 98°C for 30 sec; 19 cycles of thermocycling at 98°C for 10 s, 60°C for 30 s, 72°C for 20 s; 72°C for 5 min. 185–225 bp PAS-seq libraries were purified on 2.5% agarose gel. PAS-seq libraries were sequenced as single-end reads on NextSeq550 (Illumina).

Two biological replicates from each house mice subspecies were merged by concatenating raw FASTQ files before downstream analysis. 5’ PAS-seq adapter sequence (CATGGG) and poly(A) tails were removed using Cutadapt (v5.2^66^) and reads shorter than 10 nucleotides after trimming were discarded. Quality filtering on the remaining reads was performed using Fastp (v0.24.0^68^) with default settings.

Afterwards, reads were aligned to the mouse reference genome (mm10) using Bowtie2 (v2.5.4^62^) in single-end mode with the parameters `-local -very-sensitive -no-unal`. Mapped reads were converted to BAM format, sorted, and indexed using SAMtools (v1.8^69^). To eliminate internal priming artifacts, the 3’ end of each aligned read was identified in a strand-specific manner using BEDTools (v2.27.1^64^). For each candidate poly(A) site, 10 nucleotide genomic sequence immediately downstream of the cleavage site was extracted from the mm10 reference genome while preserving strand orientation. Candidate sites were discarded if the downstream sequence contained either six consecutive adenines or at least seven adenines within the 10-nucleotide window as these patterns can be indicative of internal priming rather than genuine polyadenylation events. The remaining reads were retained as high-confidence poly(A) sites. Strand-specific reads per million (RPM)-normalized BigWig files were used for visualization.

### Synteny analysis across house mouse subspecies

Genomic coordinates were converted between mouse genome assemblies using UCSC liftOver tool (Kent Utilities v492^72^). Genomic regions in BED12 format were first converted to BED6 format before coordinate conversion. To maximize the recovery of orthologous regions while maintaining mapping accuracy, liftOver was performed using four minimum sequence identity thresholds `-minMatch` = `0.6`, `0.5`, `0.4`, and `0.3 - minBlocks = 0.1`. Coordinates were converted from the mm10 reference genome to the CAST/EiJ genome using a custom chain file `mm10ToGCA_921999005.2.over.Cast.gz`. For the PWK/PhJ genome, coordinates were first converted from mm10 to mm39 using the UCSC chain file `mm10ToMm39.over.chain.gz` and subsequently from mm39 to the PWK genome using a custom chain file `mm39ToGCA_001624775.1.over.PWK.gz`. Mapped and unmapped regions were retained separately at each liftOver step for quality assessment. Regions successfully mapped at any of the tested `-minMatch` thresholds were combined, and duplicate entries resulting from successful mappings at multiple thresholds were removed to generate a final non-redundant set of genomic coordinates for downstream analyses.

### Sequence conservation analysis across genus *Mus* (PhastCons score)

To examine DNA sequence conservation across the genomes of genus *Mus*, we obtained whole-genome multiple sequence alignments from the UCSC Mouse Strains Assembly Hub. The alignment included the C57BL/6J (mm10) reference genome along with 18 laboratory and wild-derived mice genomes (129S1/SvImJ, A/J, AKR/J, BALB/cJ, C3H/HeJ, C57BL/6NJ, CBA/J, CAST/EiJ, CAROLI/EiJ, DBA/2J, FVB/NJ, LP/J, NOD/ShiLtJ, NZO/HlLtJ, PWK/PhJ, PAHARI/EiJ, SPRET/EiJ and WSB/EiJ). Whole-genome alignments in HAL format were converted into chromosome-specific MAF files using `hal2maf` (HAL Tools v2.2^73^) and subsequently converted into PHAST sufficient-statistics (SS) format using `msa_view` (PHAST v1.6^74^). We next performed neutral evolutionary model estimated from putatively neutral genomic regions. To define neutral genomic regions, protein-coding genes, coding sequences (CDS), untranslated regions (UTRs), promoters, lincRNA genes, and ENCODE candidate cis-regulatory elements (cCREs), including promoter-like and enhancer-like elements, were masked using GENCODE M25 and ENCODE annotations. Fixed-length windows were randomly sampled from the remaining non-functional regions across the genome and converted into SS alignments for model training. A neutral substitution model was then estimated using `phyloFit` (PHAST v1.6^74^) under the HKY85 substitution model with expectation-maximization and high numerical precision while keeping the phylogenetic topology fixed.

The resulting neutral model was used to calculate genome-wide conservation scores. PhastCons was used to estimate the probability that each nucleotide belongs to a conserved element. Chromosome-specific conservation scores were generated in WIG format and converted to BigWig format using `wigToBigWig` (UCSC Kent Utilities v492^72^). Feature-level conservation scores were subsequently extracted from the BigWig files using `bigWigAverageOverBed` (UCSC Kent Utilities v492^72^).

### Sequence conservation analysis across placental mammals (PhastCons score)

To evaluate sequence conservation over a long evolutionary timescale, we used publicly available conservation tracks from the UCSC Genome Browser. We downloaded the PhastCons BigWig file (mm10.60way.phastCons60wayPlacental.bw) corresponding to the placental mammal subset of the UCSC 60-way vertebrate multiple alignment. These conservation tracks were generated using the mm10 (GRCm38) mouse reference genome together with 39 additional placental mammalian genomes (40 species in total). Conservation scores were extracted from the downloaded BigWig files using `bigWigAverageOverBed` (UCSC Kent Utilities v492^72^). The same genomic annotations and feature definitions described above were used throughout the analysis.

### Synteny analysis across placental and non-placental mammals

Mouse pachytene piRNA genes overlifted to the corresponding genome assemblies of human (hg19), rhesus macaque (rheMac8), marmoset (calJac3), rat (rn6), opossum (monDom5), and platypus (ornAna1) using the UCSC liftOver tool (Kent Utilities v492^72^). Genomic regions in BED12 format were first converted to BED6 format before coordinate conversion. Given that the analysis involved evolutionarily divergent species, liftOver was performed using a relaxed minimum match threshold `-minMatch=0.1`.

Successfully mapped and unmapped regions were retained separately for quality assessment. For each species, the lifted coordinates were compared with previously annotated piRNA loci^12^ using a strand-aware overlap analysis. Lifted regions with a best overlap fraction ≥50% were classified as evolutionarily conserved.

### Statistical analysis

Statistical analyses and graph generation were conducted using R v4.3.2 (https://www.rstudio.com). Box plots were used to present data distribution. Boxes represent interquartile ranges (IQRs), spanning the first and third quartiles. Outliers were defined as data points with values > third quartile + 1.5 × IQR or lower than the first quartile − 1.5 × IQR, where IQR is the difference between the maximum of the third and the minimum of the first quartile. Significance was measured using two-sided unpaired Student *t* test in Figure 2b, 2d and 3a. Relationship between variables were measured using spearman’s correlation (P) in Figure 3c. Significance was measured by two-sided wilcoxon matched-pairs signed rank sum test in Figure 4d and Extended Data Figure 6d and 7d. DESeq2 analysis with Benjamini-Hochberg correction was used in Figure 5b. Fisher’s exact test was used to calculate *p* values in Figure 5c. Kruskal-Wallis test, pairwise Wilcoxon–Mann-Whitney test with Benjamini-Hochberg correction, adjusted *p* values was used in Extended Data Figure 8b, 8c, 8e and 8f.

## Data and code availability

Sequencing data are available from the National Center for Biotechnology Information Sequence Read Archive using Gene Expression Omnibus (GEO) accession number GSE336725.

## Acknowledgments

We thank the personnel of Stockholm University animal facility with a particular gratitude to S. Olsson, S. Oerther, and R. Askar for their excellence in mouse colony management; Matthew Hunt for proofreading English grammar and clarity; National Genomics Infrastructure – funded by Science for Life Laboratory, the Knut and Alice Wallenberg Foundation and the Swedish Research Council, and SNIC/Uppsala Multidisciplinary Center for assistance with massively parallel sequencing and access to the UPPMAX computational infrastructure. This work was supported by the Swedish Research Council grant 2020-03818 (D.M.Ö), the Swedish Research Council grant 2024-04321 (D.M.Ö), Carltryggersstiftelse CTS 21:1158 (D.M.Ö), and Åke Wibergs Stiftelse M23-0022 (D.M.Ö).

## Author contributions

M.S., J.L.F., Y.T.X., and D.M.Ö. conceived and designed the experiments. M.S., M.A., A.E., and M.M.A. performed the experiments. M.A., and Y.T.X. analyzed the sequencing data. M.S., Y.T.X., and D.M.Ö. wrote the manuscript. All authors discussed and approved the manuscript.

