## Supplementary figures and images for "Evolutionarily labile pachytene piRNAs target an altered set of mRNAs in male hybrids of house mouse subspecies"

### Extended Data Figure 1

Säflund et al., Extended Data Figure 1

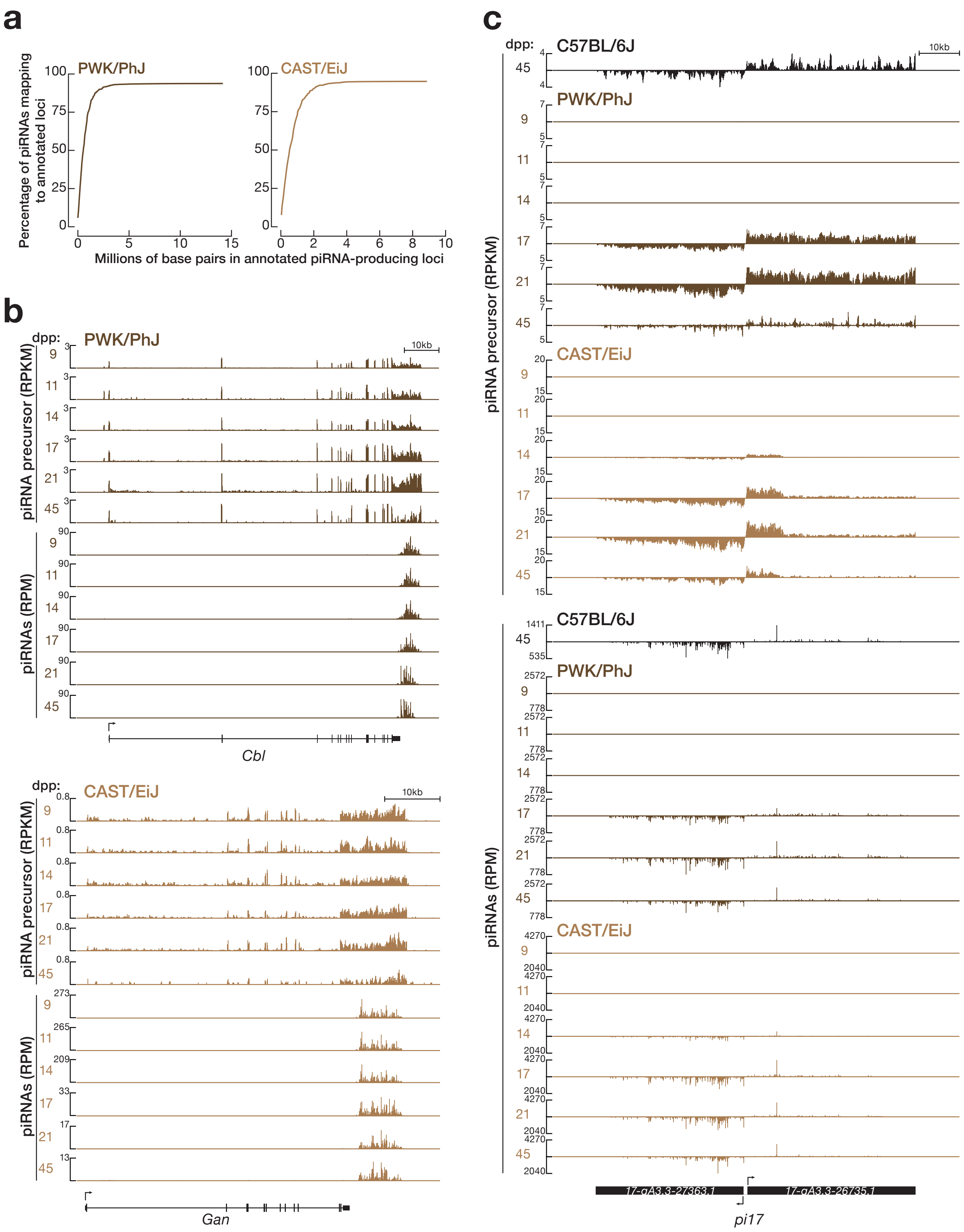

### Extended Data Figure 2

Säflund et al., Extended Data Figure 2

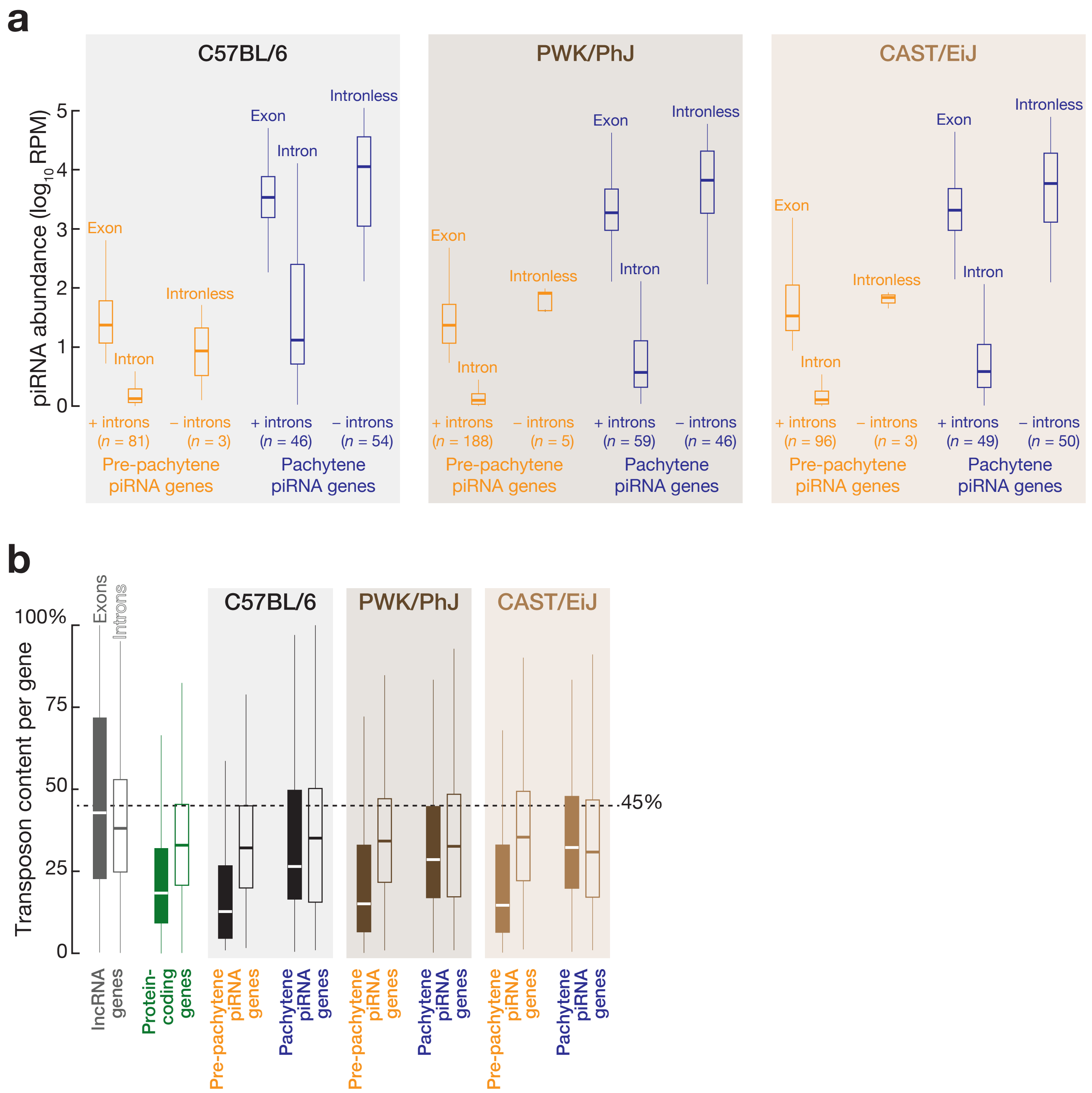

### Extended Data Figure 3

Säflund et al., Extended Data Figure 3

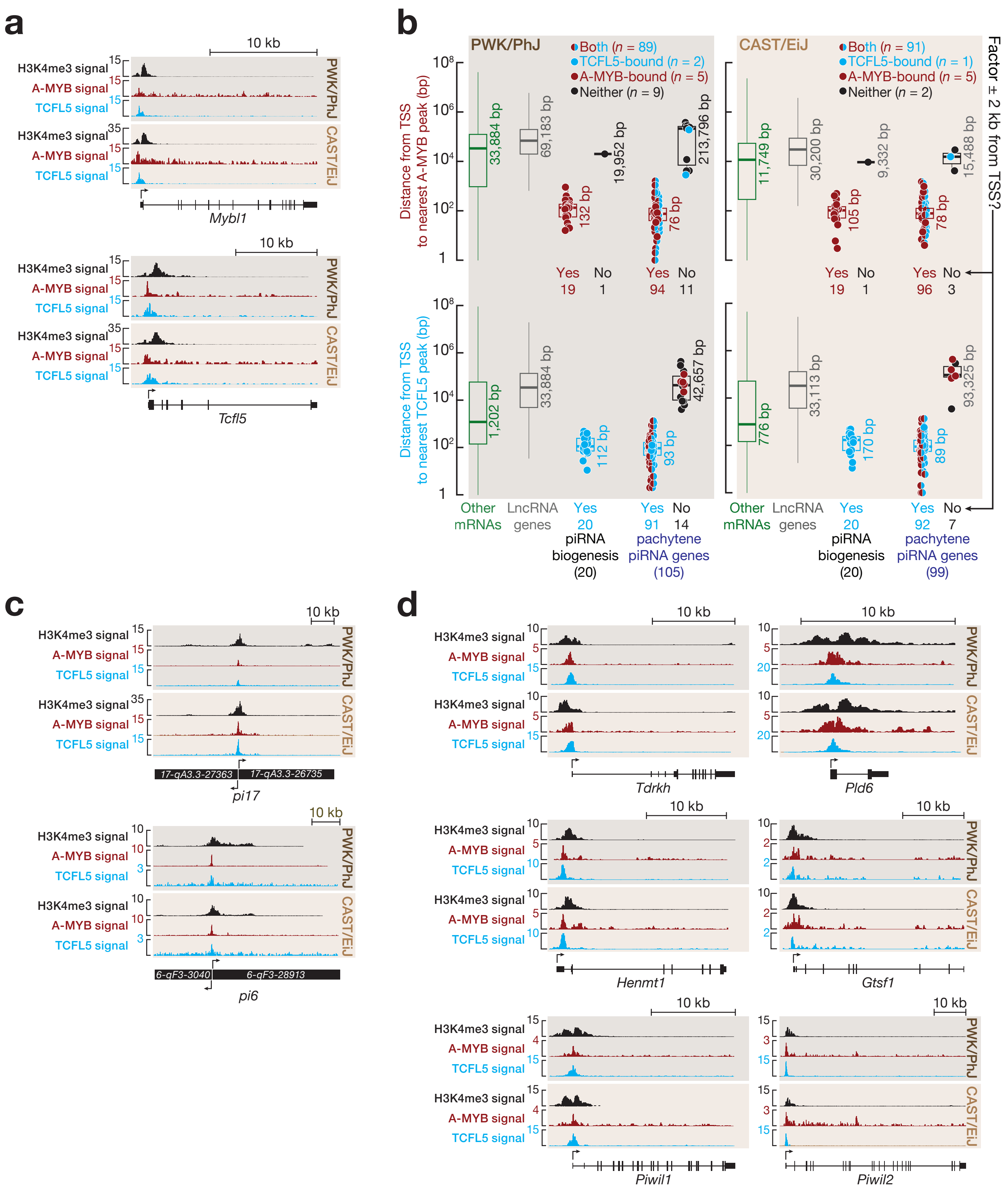

### Extended Data Figure 4

Säflund et al., Extended Data Figure 4

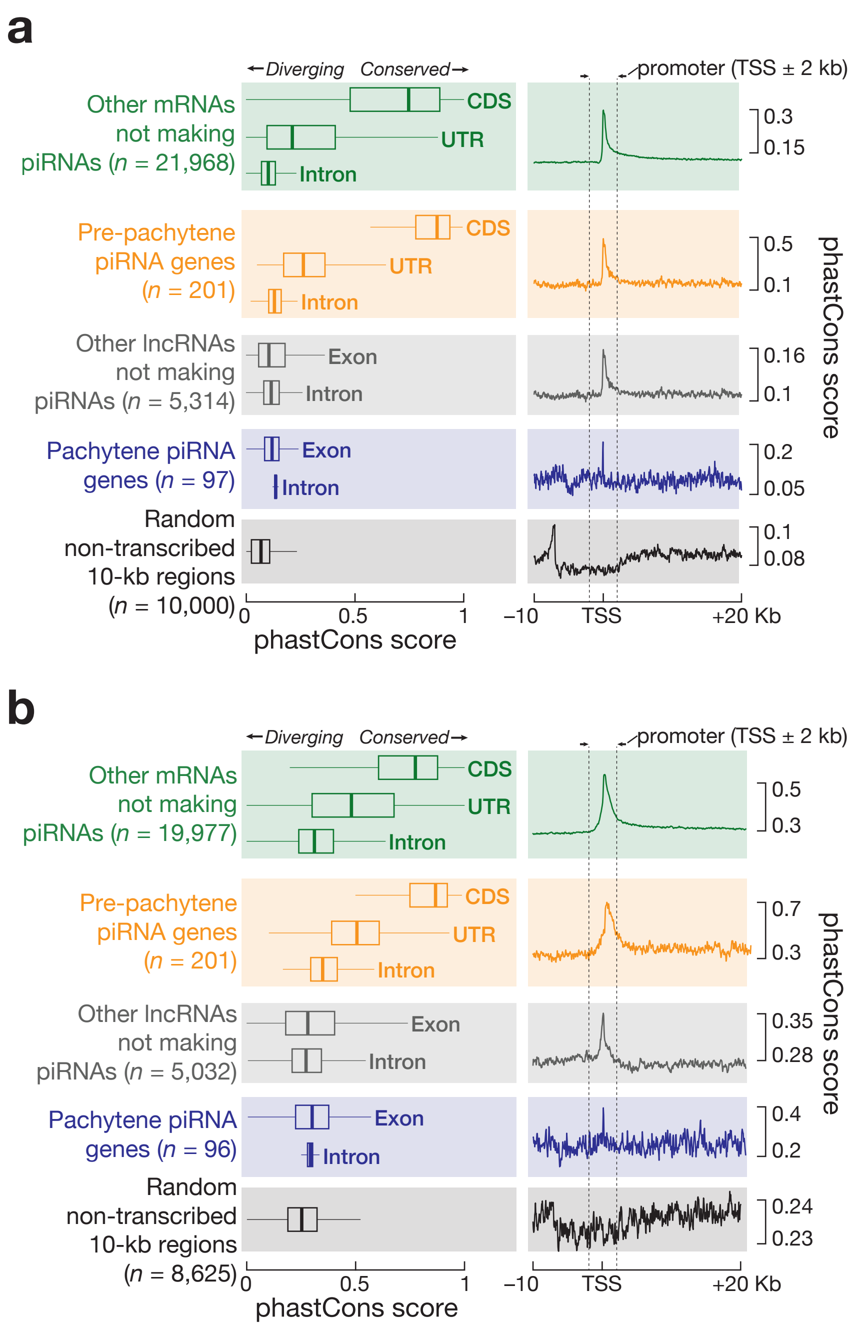

### Extended Data Figure 5

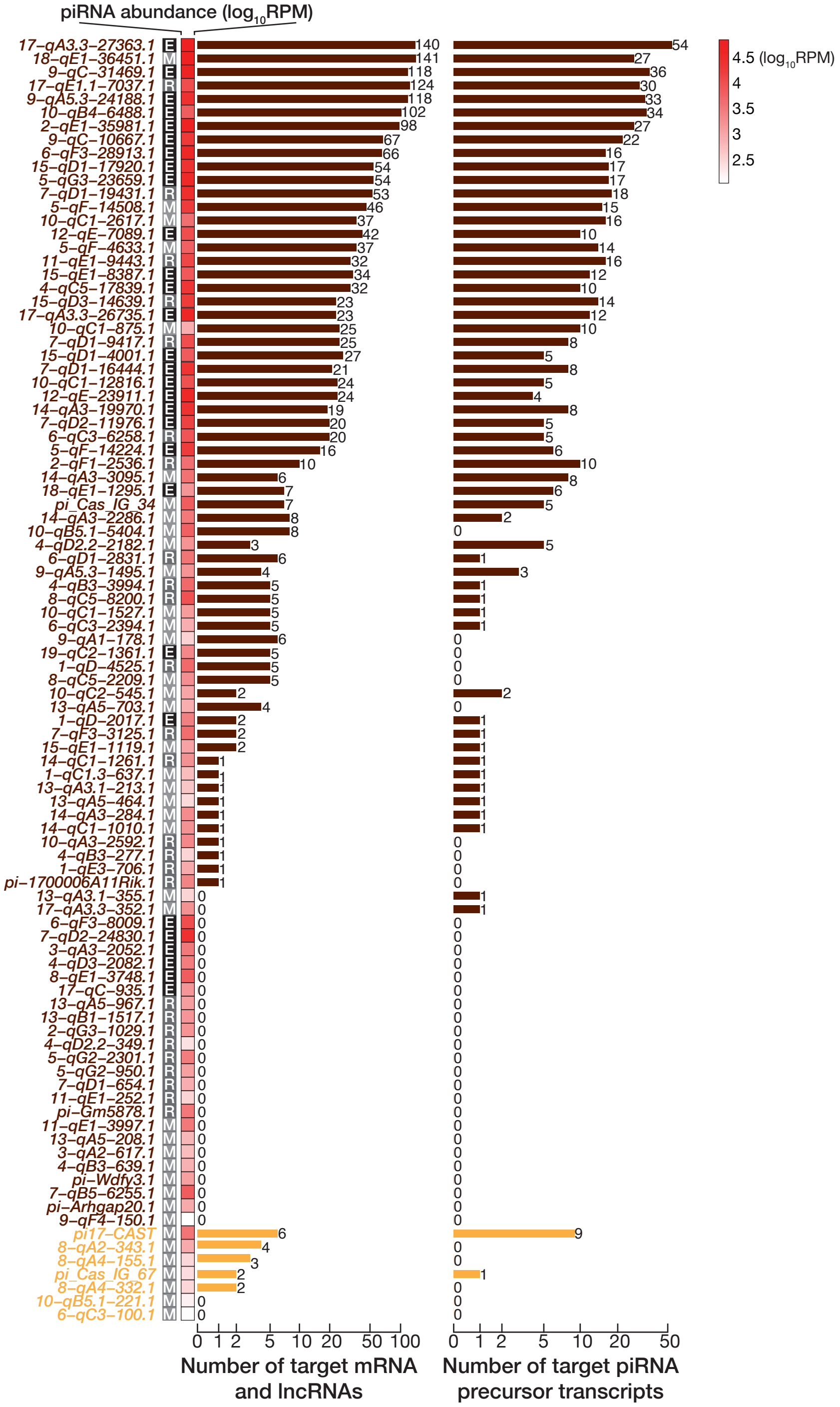

### Extended Data Figure 6

Säflund et al., Extended Data Figure 6

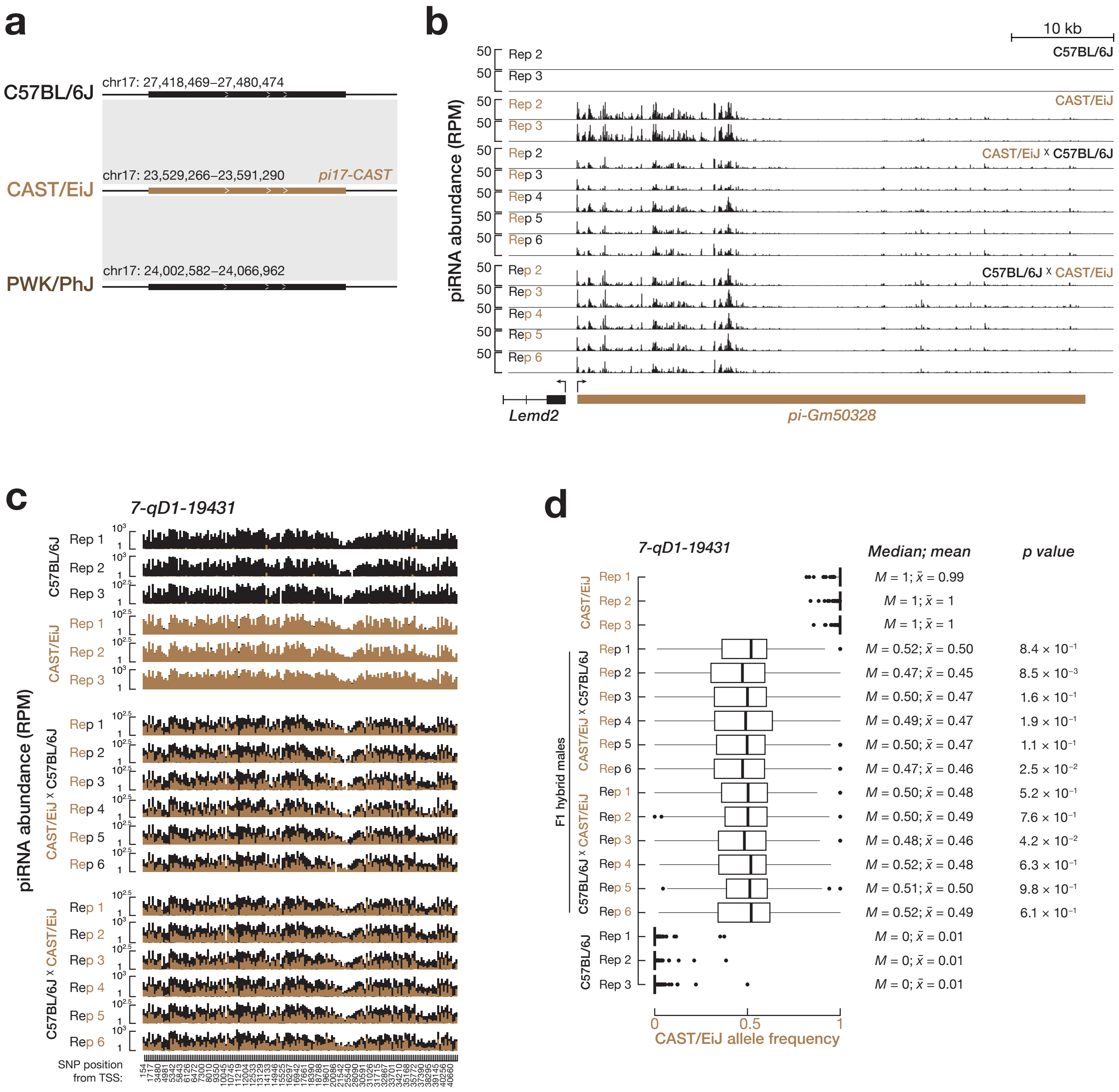

### Extended Data Figure 7

Säflund et al., Extended Data Figure 7

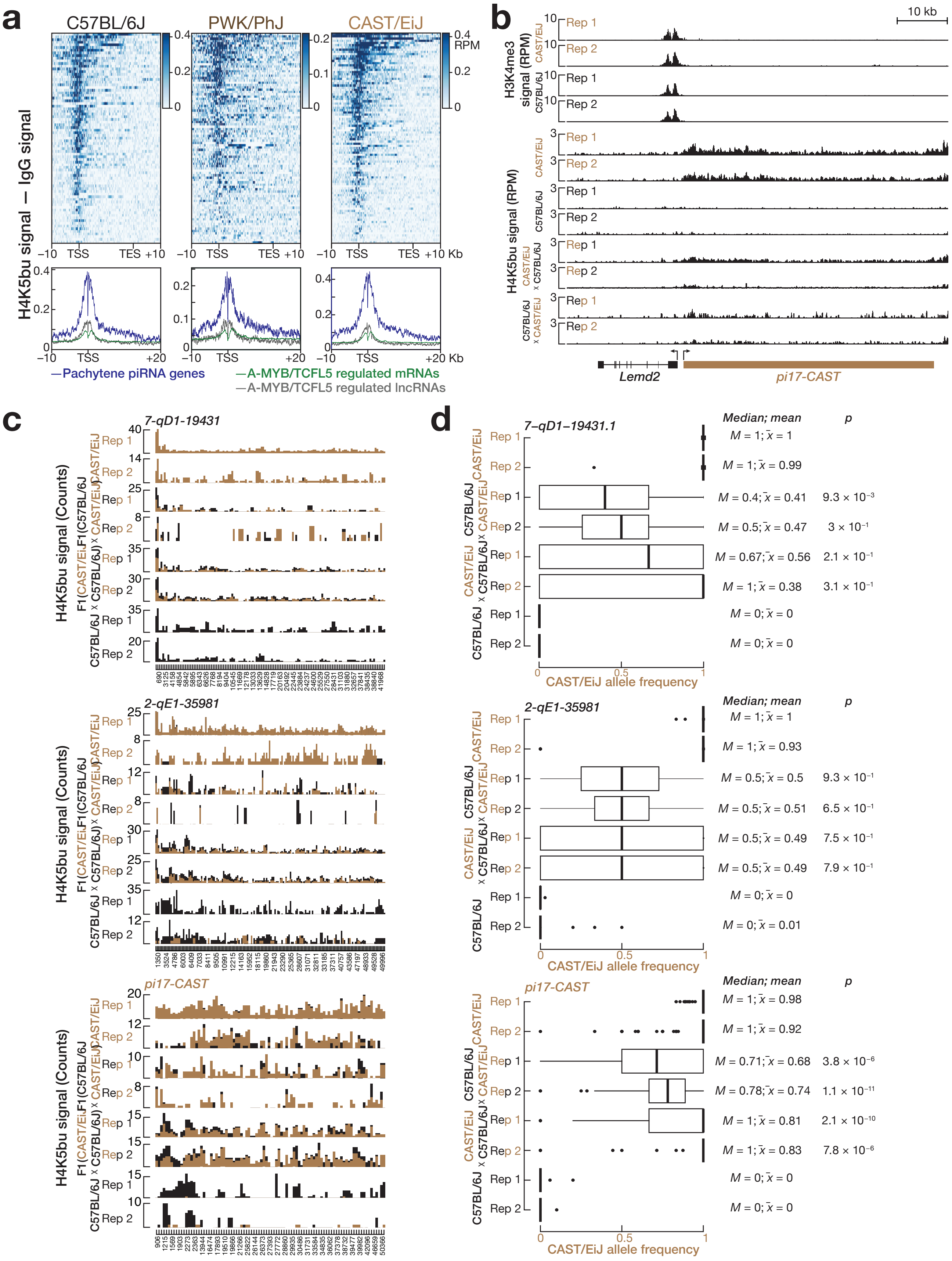

### Extended Data Figure 8

Säflund et al., Extended Data Figure 8

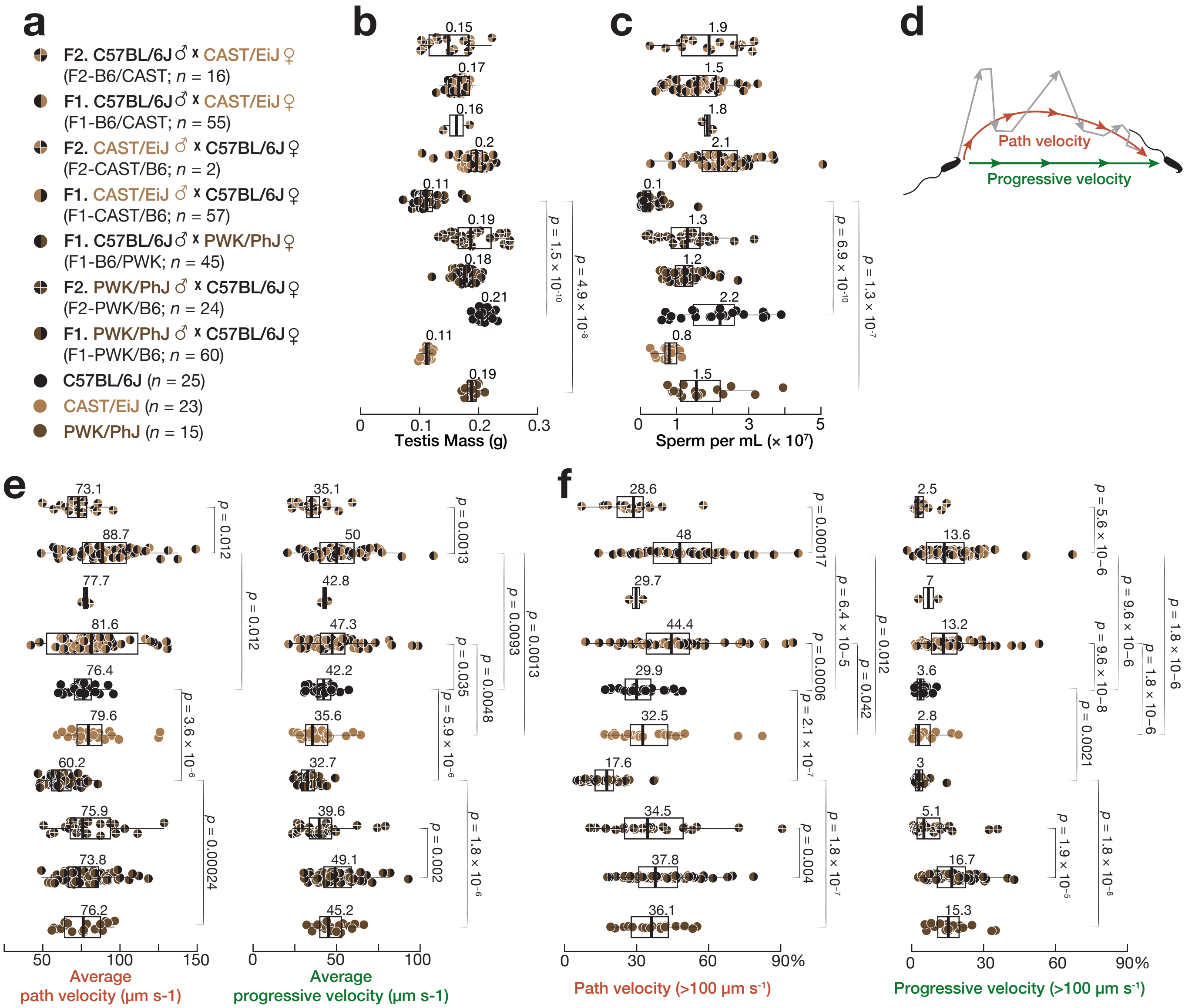
